# Antimicrobial resistance in a warmer world: selection and transfer of kanamycin resistance is temperature dependent in *E. coli*

**DOI:** 10.64898/2026.10.16.759609

**Authors:** Zoltan Erdos, Tatiana Dimitriu, Ben Raymond, Angus Buckling, Daniel Padfield

## Abstract

Climate warming consistently correlates with increased antimicrobial resistance (AMR). Understanding the biological mechanisms underpinning this relationship is important for both prediction and mitigation of the future burden of antibiotic resistance in a warmer world, especially given AMR can spread through both selection and horizontal gene transfer. Here, we experimentally measure how temperature alters susceptibility to kanamycin, selection and spread of resistance in *Escherichia coli* MG1655 carrying the natural plasmids R1 and RP4. We predict that susceptibility should be lowest - and plasmid transfer highest - at maximal growth rate. Thermal performance of bacterial growth peaked at 40°C and we detected a small plasmid cost - for RP4, but not R1 - in the absence of kanamycin. The thermal performance of plasmid transfer did not mirror that of bacterial growth, instead, R1 transfer peaked at 35°C with a much narrower thermal tolerance. Susceptibility to kanamycin decreased with increasing growth rate until 31°C, before increasing again as temperatures approached and exceeded optimal growth temperature. Increasing susceptibility coupled with high growth rates meant that selection was strongest at the highest measured temperature, but the shape of the relationship changed with kanamycin concentration. In mixed populations of resistant and susceptible bacteria, we found that AMR spread most where selection was strongest, irrespective of plasmid transfer. Our findings suggest increased AMR at higher temperatures is predominantly driven by higher growth rates and increased susceptibility, resulting in stronger selection. Our study provides a framework to uncover generalities across antibiotic classes, strains, species, and plasmids, laying the foundation for improved prediction of AMR selection in the environment.

## Introduction

Recent studies have found a positive association between environmental temperatures and levels of antimicrobial resistance (AMR) in environmental and clinical bacteria (1–4). While these studies are generally correlational, field experiments have also identified a positive relationship (5,6), suggestive of temperature-driven selection of resistance. However, there remains little experimental data examining the biological mechanisms underpinning the relationship between temperature and AMR. This knowledge gap limits our ability to understand, predict, and ultimately mitigate its future burden.

Temperature-driven selection for resistance could increase three key drivers of AMR: rates of *de novo* evolution of resistance, relative abundance of already resistant bacteria, and horizontal gene transfer. This stronger selection arises from the fundamental impact temperature and antibiotics have on biological processes, particularly growth rate, and how the two interact. As temperatures increase, growth rates generally increase up to an optimum, before rapidly declining. This relationship can be described using thermal performance curves (TPCs), which are summarised by several key “traits” (e.g. optimum temperature, maximum rate) that characterise the shape of the curve. These traits can, in turn, be used to describe thermal adaptation (7,8) and predict responses to climate change (9). Antibiotics also strongly influence growth rate, while the dose response of bacteria to antibiotics can be characterised by quantitative “traits” such as the no significant effect concentration (NSEC)(10), half maximal effective concentration (EC_50_), and minimum inhibitory concentration (MIC)(11). These measures summarise antibiotic susceptibility and selection at low to lethal concentrations. Despite extensive work on the impacts of temperature and antibiotics on bacterial growth, their joint effect remains poorly understood.

Temperature alters the impact of antibiotics on susceptible bacteria, such that the same concentration can have a different effect across temperatures (12). In this way, antibiotics shift the thermal performance of *E. coli* in the short-term (13) and as a result of evolution (14,15), while temperature likewise shifts antibiotic performance both in the short term and through the evolution of resistance (16,17). However, no studies have investigated how temperature alters the antibiotic dose-response, a key determinant of antibiotic selection, but we can make some simple predictions. For example, if antibiotic susceptibility (defined by NSEC, EC_50_, and MIC) does not change across temperatures, selection should be strongest at optimum temperatures (maximum drug-free growth rate). However, susceptibility might be growth-dependent, and thus also change with temperature. Recent mechanistic work that modulated growth rate using nutrients found that faster growing cells were less susceptible to kanamycin (measured as EC_50_) (19). If this holds for temperature, susceptibility will be lowest at optimal temperature, with the temperature-response of selection emerging from the balance between how temperature changes susceptibility and drug-free growth rate.

Temperature could further influence antibiotic selection by altering the fitness cost of resistance. Antibiotic resistance is typically assumed to be costly, with susceptible bacteria outcompeting resistant strains in the absence of antibiotics (20,21). Yet, these costs vary across environments and strains (22–24), and resistance genes that are costly at one temperature can be neutral or beneficial at another (25). Because of this complexity, we have no *a priori* expectation of how costs of resistance will vary across temperature or in relation to a TPC. However, if costs of resistance are minimised at higher growth rates, then this would also increase antibiotic selection at optimal growth temperature.

Irrespective of selection, higher temperatures could result in increased spread of AMR through horizontal gene transfer, particularly via plasmids (26). Theory and experimental work suggest that plasmid transfer increases at high host growth rates (27–29), such that it would be highest at the optimal temperatures of the donor and recipient. This prediction becomes more difficult when there are thermal mismatches - differences in the shapes of the TPCs - between the donor and recipient. In this situation, we expect plasmid transfer to only occur at temperatures where the thermal tolerance of hosts overlap and where both the donor and recipient have positive growth rates. Temperature can also have a plasmid-dependent effect on transfer independent of its effect on host growth rate. For example, IncHI1 plasmids have increased transfer rates in a range of 22-30°C (30), even though their predominant hosts - *Salmonella* spp. - grow best at temperatures close to 37°C (31). The nucleoid protein histone-like nucleoid-structuring protein (H-NS) represses conjugation of IncHI1 plasmids at temperatures above 30°C (32), while it upregulates transfer of IncX3 plasmids in warm temperatures (33). This suggests that plasmids themselves are adapted to transfer at certain temperatures, but the relative importance of plasmid thermal preference versus host growth rate for transfer is untested.

Here, we experimentally tested how temperature alters the selection and spread of AMR using the antibiotic kanamycin in the model organism, *Escherichia coli* MG1655, the IncFII plasmid R1, and the IncP plasmid RP4. We predict that selection for kanamycin resistance will change non-linearly with temperature, with selection strongest at optimum temperatures, but this could change depending on the strength of the relationship between susceptibility and temperature. Regarding plasmid transfer, we expect it to peak at optimum host growth temperature and only occur when the donor and recipient have positive growth rates. To test these predictions, we measured selection for resistance by quantifying growth of resistant (plasmid carrier) and susceptible bacteria across 10 antibiotic concentrations at 11 temperatures. Separately, we quantified plasmid transfer at these same temperatures in the absence of antibiotics, and conducted invasion experiments to determine how temperature alters AMR selection in mixed populations of resistant and susceptible bacteria.

## Methods

### Bacterial strains, plasmids, media and growth conditions

*E. coli* MG1655 (wt) and *E. coli* MG1655 carrying plasmid R1 or RP4 were used in this study. A spontaneous nalidixic acid-resistant mutant (nal^R^) was derived from the wt strain as the recipient in conjugation assays. To derive nalidixic acid-resistant mutants, a single colony of *E. coli* MG1655 was inoculated into 5 mL of antibiotic-free Luria–Bertani (LB) broth and incubated at 37°C with shaking at 180 r.p.m for 24 hours (h). The culture was centrifuged at 13,000 × g for 4 minutes (min), the supernatant discarded, and the pellet resuspended in sterile water. This washing step was repeated before the cells were resuspended in 100 μL of 8.5 g/L saline and spread onto LB agar plates supplemented with nalidixic acid (64 μg/mL). Plates were incubated at 37°C for 24h. Emerging mutant colonies were subcultured every 24h for three days on LB agar supplemented with nalidixic acid (64 μg/mL) to confirm stable resistance. Selected mutants were stored in glycerol stocks (final concentration 25%) at −70°C. The two naturally occurring conjugative plasmids used were: R1, a narrow host-range IncFII plasmid; and RP4, a broad host-range IncP plasmid. Both plasmids confer resistance to antibiotics, including kanamycin via genes APH(3’)-Ia and APH(3’)-Ib, respectively.. Both plasmids were introduced into the wt strain by conjugation from an *E. coli* MFD*pir* donor strain (34).

For routine propagation and assays, strains were streaked onto LB agar supplemented with the appropriate selective antibiotic (50 μg/mL kanamycin for R1 and RP4; 64 μg/mL nalidixic acid for nal^R^), incubated at 37°C for 18h, and stored at 4°C until required. Liquid overnight cultures were prepared by inoculating a single colony into 5 mL LB broth containing the respective antibiotic (except for wt) and incubating at 37°C with shaking at 180 r.p.m. Prior to use in assays, overnight cultures were diluted 20-fold in fresh LB.

### Measuring dose response across temperatures

To characterise how temperature and antibiotic concentration jointly affect bacterial growth, dose-response assays were performed by broth microdilution in 96-well plates (11). A 1.5-fold serial dilution of kanamycin in LB was prepared over a range of 0–64 μg/mL (10 different concentrations), accounting for the 5 μL inoculum volume. Each column of a 96-well plate was loaded with 195 µL of a single kanamycin concentration, with negative controls consisting of blank wells containing 200 µL sterile LB. Each well was then inoculated with 5 μL of 20-fold diluted overnight culture of wt, wt + R1, wt + RP4 or nal^R^ and 50 µL mineral oil was overlaid onto each well to minimise evaporation. Optical density at 600 nm (OD_600_) was monitored at 11 temperatures ranging from 16–45°C on a Multiskan SkyHigh microplate spectrophotometer (Thermo Fisher Scientific) with shaking for at least 24h, but up to 99h at slower growing temperatures. Each temperature was run at least twice, yielding at least four replicates per strain per temperature–dose combination. Some methods to estimate conjugation rate require using growth rates of donors, recipients and transconjugants, so the growth rate of transconjugants (nal^R^ + R1, nal^R^ +RP4) were measured in the absence of antibiotics using the same protocol and temperatures as described above.

### Plasmid transfer assays

To characterise how temperature affects plasmid transfer, conjugation assays were performed across a temperature range of 13–46°C using wt carrying R1 or RP4 as the donor and nal^R^ as the recipient. Overnight cultures of each strain were prepared in 5 mL LB containing the appropriate antibiotic. Prior to conjugation, overnight cultures were diluted 10-fold into fresh LB without antibiotics and incubated at 37°C for 1h to reach exponential phase, then pelleted by centrifugation (8000 x g, 2 min) and resuspended in fresh LB. Conjugation assays were performed in 96-well plates pre-loaded with 80 μL LB. Donor and recipient cultures were mixed in equal volumes (10 μL each) and mated for 3 h (100 μL total volume) in a VeritiPro 96-Well Thermal Cycler with a 6-zone Veriflex Block (Applied Biosystems), enabling temperature control of six zones within a single plate. Using two thermal cyclers simultaneously, matings were conducted from 13-46°C at 3°C increments from the same set of overnight cultures. The entire experiment was repeated seven times. Donor, recipient, and transconjugant densities were estimated by serial dilution and plating onto selective media: plasmid-bearing cells were enumerated on LB agar supplemented with kanamycin (50 μg/mL), recipients on LB agar supplemented with nalidixic acid (64 μg/mL), and transconjugants on double-selective plates containing both antibiotics. Each assay included donor-only and recipient-only control wells, which showed no growth on double-selective plates.

Conjugation rates (γ) were estimated as γ = T / (D · R · t) (mL cell⁻¹ h⁻¹), where T, D, and R are the densities of transconjugants, donors, and recipients (cells mL⁻¹), respectively, and t is the incubation time (h). To check the suitability of this estimation method, we compared it to those derived using the Approximate Extended Simonsen (ASM) model using the R package ‘*conjugator*’ (35). The ASM model accounts for population specific growth rates (*ψD*, *ψR*, *ψT*). There was a strong correlation between the two methods (Figure S1), so we used the simpler method described above for the analyses.

### Competition (invasion-from-rare) assays

To test whether temperature-dependent differences in growth rate translate into selection for resistance in mixed populations, competition assays were performed across a range of kanamycin concentrations. Mixed inocula comprising 10% donor (wt + R1 or wt + RP4) and 90% nal^R^ (20 μL total) were added to 120 μL LB containing kanamycin at concentrations of 0, 2, 3.5, 5, or 15 μg/mL in 96-well plates. Cultures were incubated at 21, 31, and 40°C in a VeritiPro 96-Well Thermal Cycler with a 6-zone Veriflex Block. Incubation times were adjusted to account for temperature-dependent differences in growth rate: 22h at 21°C, 4h at 31°C, and 3h at 40°C. Each temperature X antibiotic combination was assayed from a single run of the thermal cycler with six technical replicates, and the experiment was repeated three times. At the end of incubation, strain densities were determined by serial dilution and plating onto selective media as described for plasmid transfer assays.

### Statistical analyses

#### Overview of open software used and modelling approach

All analyses used R (v4.5.2) (36) and scripts and data are available on GitHub (https://github.com/padpadpadpad/ecoli_temperature_amr). All R scripts used the suite of packages known as the ‘*tidyverse*’ (37). We fitted nonlinear models in a Bayesian framework to characterise (a) the thermal performance curves of the strains in the absence of antibiotics and (b) dose-response curves of *E. coli* MG1655 wild-type and nal^R^ with and without the plasmids R1 and RP4 across temperatures. Bayesian models were fitted using ‘*brms*’ (38) and were processed using ‘*tidybayes*’ (39). We employed a small model implementation approach where we achieved convergence of a least complex model on a reduced dataset before adding complexity. All models ran for at least 3000 iterations with 4 chains. Weakly informative priors were used to facilitate model convergence, which was assessed using comparisons of the prior and posterior density distributions (Figures S3–S14), Rhat values (all values were <1.01) and manually checking chain mixing. Summaries of predictions and parameters present estimations of 95% highest density credible intervals.

#### Processing growth rate assays

For each well, we estimated exponential growth rate. We corrected each well by the plate-specific median blank value over the course of each run, after checking that no blank wells had been contaminated. We used ‘*gcplyr*’ (40) to calculate growth metrics. Exponential growth rate was estimated by taking the maximum per capita derivative of the OD measurements across every 60 minute window (representing 6 measurements) using ‘*calc_deriv()*’. A set of steps were then used to clean the growth rate estimates for the dose-response assays. Of 2640 estimates of growth rate, 95.8% passed quality control and were retained.

First, we replaced any missing estimates with 0 because ‘*calc_deriv()*’ likely failed due to low insufficient growth in those wells. If the maximum density was zero, we classified *gcplyr* as failing. If >50% of the blank-corrected OD in any well was below 0.0090 (representing a very low bacterial density), we set the estimates of growth rate to 0 as there was very little growth. Wells that dried up during the assay (likely because mineral oil did not cover the whole surface) were identified when final density was less than 60% of the maximum density, and where maximum density is greater than 0.075, because dried wells are harder to identify when there is little growth. Some wells at high antibiotic concentrations had zero growth for over a day before rapidly increasing, a hallmark of *de novo* evolution of resistance. To identify these, we grouped wells across temperatures and by whether the strains were resistant or susceptible and looked at variation in each group’s final corrected OD reading by running ‘*performance::check_outliers()*’ with ‘*method = ‘zscore_robust’*’ to identify potential outliers (41). We classified outliers which had a final OD more than 0.15 over the group median as suspected *de novo* evolution, and any outliers that were 0.5 OD or more below the median as failed wells. We compared our programmatic way of identifying outliers with those identified as failed during the assays. Any wells where classification differed was checked by two of the authors (Padfield and Erdos) independently. If there was agreement between the authors on the growth rate estimate being fine, then the estimate was kept, if not then the well was classified as failed. After this quality control, we removed 13 wells because they had failed, 29 because they had dried up, and 69 because we suspected *de novo* evolution (an example plate check is Figure S2). The pipeline is included with the data and scripts used (see *05_clean_data.R*), and we believe represents a transparent approach to data processing that combines automation with domain knowledge. For the growth responses of nal^R^ + R1 and nal^R^ + RP4 - which were required to estimate transconjugant growth rates - no cleaning was needed, but we visually checked all the curves.

#### Analysing thermal performance curves

We fitted thermal performance curves using a modified version of the *flexTPC* (42) equation because of its flexibility and its biologically meaningful parameters that made the selection of custom priors easier. The equation is:

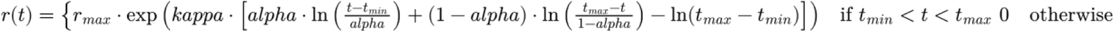

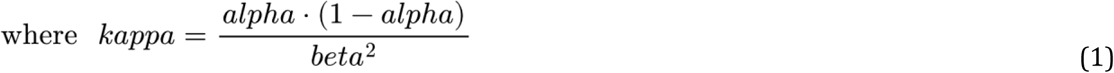

Where *r(t)* is the rate at any given temperature in °C (t), *r_max_* is the maximum rate, *t_min_* is the minimum temperature at which the rate is 0 and *t_max_* is the maximum temperature at which the rate is 0. *alpha* (bounded between 0 and 1) represents the skewness of the curve, where 0 corresponds to optimum temperature being equal to *tmin*, 1 to optimum temperature being equal to *t_max_*, and 0.5 being a symmetrical curve where the optimum temperature, *t_opt_*, is equal to (*t_min_* + *t_max_*)/2. *beta* determines the breadth of the thermal performance curve near its peak, with larger values corresponding to broader curves and smaller values to narrower curves. Optimum temperature (*t_opt_*) can be calculated as:

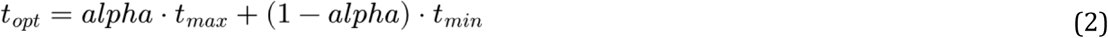

To improve model convergence, we modified the equation to allow a more gradual transition from zero to positive rates inside *t_min_* and *t_max_*, and transformed parameters so that they were of similar magnitude and constrained in terms of their sign. All modifications of the model structure were checked to ensure estimated parameters had the same meaning and formulations were equivalent (see *00_flextpc_check.R*).

We fitted thermal performance curves to estimates of drug-free growth rate for all strains by plasmid combinations using Equation 1 in a Bayesian framework. The drug-free growth rate is important for antibiotic selection as it sets the height of the dose response curve. For example, if a bacterium is very susceptible, but is at a temperature under extreme growth-limitation, absolute selection will be small because there is limited scope for growth at that temperature. The total number of data points fitted to this model was 933. For each parameter, we included a fixed effect of strain (wt, wt + R1, wt + RP4, nal^R^, nal^R^ + R1 and nal^R^ + RP4). Models were run with a gaussian error structure. The curves for the nalidixic acid mutant were modelled to estimate recipient (nal^R^) and transconjugant growth rates (nal^R^ + plasmid) for calculating plasmid transfer rates using the Approximate Extended Simonsen (ASM) model. The code for fitting TPCs can be found in *brms_tpc.R*, and priors can be found in Table S1.

To model horizontal gene transfer, we fitted thermal performance curves separately to the transfer rate of R1 and RP4. The total number of data points included in these models were 96 and 84 respectively. We included a random effect of replicate (representing a different overnight) on each parameter, which were marginalised when calculating the overall plasmid transfer rate (e.g. predictions included the random effects and then the mean was taken). Models were run with a gaussian error structure. The code for fitting this model can be found in *brms_plasmid.R*, and priors can be found in Table S2. For all thermal performance curves, we calculated the key traits of optimum temperature, maximum growth rate, cardinal temperatures (*t_max_* and *t_min_*), and thermal tolerance (*t_max_* -*t_min_*).

#### Analysing dose-response curves

We fitted dose response curves to the growth rates of wild-type, nalidixic acid mutant and the plasmid carriers using the *ecxwb1* model from the R package *‘bayesnec*’ (43). This is a 4-parameter sigmoidal decay model, given by the equation:

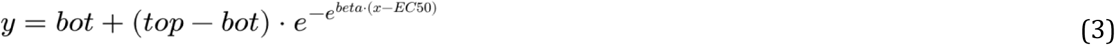

Where *bot* represents the lower plateau for the response at infinite concentration of the predictor, *x*, *top* is the y-intercept at zero concentration, *beta* is the exponential decay rate of the response, and *EC_50_* is the effective dose where the rate is 50% of top. This model was chosen over the more commonly seen log-logistic models as it provided more flexibility in the curve shape by not being symmetrical around *EC_50_* on the transformed scale. The total number of data points fitted to this model was 2522. To ensure even spacing of the data across kanamycin concentrations, and to include the 0 µg/mL data, we used an inverse hyperbolic sine transformation on the kanamycin concentrations. To improve model convergence, we transformed growth rate to be a proportion of maximum growth rate by dividing each growth rate by the highest observed in the dataset (44). If any values were equal to 0 or 1 after this transformation, we added or subtracted 10^-6^ respectively. We used the beta family and default priors from *bayesnec* for *beta*, *bot*, and *top*, and an *EC_50_* prior based on the mean concentration of kanamycin on the transformed scale. For each parameter, we included a random effect for each unique combination of temperature and strain to allow parameters to vary across individual curves. The full model included the nal^R^ data because - despite not being used in downstream analyses - it increased the number of levels available for partial pooling.

#### Analysing changes in AMR selection

For the dose response of the wild-type bacteria, we estimated five traits that are important for the selection and evolution of resistance: the no-significant effect concentration (NSEC), the concentration at which the growth rate is reduced by half (EC_50_) the minimum inhibitory concentration (MIC), and the sub-MIC selection window. The NSEC is conceptually similar to the traditional no-observed-effect concentration and no-effect concentration but it decouples the estimate from being directly dependent on the placement of treatment concentrations and requires statements of precision (10). It is calculated as the concentration at which the growth rate is different to the 95% credible interval at zero concentration for each individual curve. This is widely interpreted as an estimate of the lowest concentration at which selection occurs. We defined the MIC as the concentration at which growth rate was 95% reduced compared to the maximum growth rate observed in the whole dataset. The sub-MIC selection window (MIC - NSEC) represents the concentrations at which kanamycin reduces bacterial growth without total inhibition. A smaller value reflects a steeper dose response curve, which should - in the absence of existing resistance - result in weaker selection for resistance evolution (45,46). These traits influence selection regardless of the cost of resistance which can differ across genotypes and environments (25,47). Code for the analysis of the dose response curves and downstream models can be found in *brms_drc.R*, and priors can be found in Table S3.

To determine how temperature changed the impact of kanamycin across concentrations, we fitted a model to the NSEC, EC_50_, and MIC, and sub-MIC selection window values estimated for each dose response curve. To propagate uncertainty, we fitted separate models to 500 independent draws and combined the posterior distributions. The response of the model were the estimates of the traits, the predictors were temperature (included as a second order polynomial) and trait (NSEC, EC_50_, MIC, or sub-MIC selection window) and their interaction. The model was fitted with a truncated normal distribution with a lower bound of 0 to prevent negative concentrations. To aid model convergence, we set a weakly informative prior on sigma based on the model fit to a normal distribution (Table S4). We then calculated where the lowest and highest values of each trait occurred to determine where selection was weakest and strongest and calculated the fold-change in that trait across temperatures.

We used predictions from the dose-response curves to estimate how selection for kanamycin resistance changed across combinations of temperature and kanamycin concentration. Selection was approximated using two metrics: the selection rate constant (*r*) and selection coefficients (*s*). The selection rate constant, *r*, was calculated as the difference between the Malthusian growth parameters of the plasmid carrying (resistant) and wild type *E.coli*, providing an estimate of the fitness effect of plasmid carriage at each assay temperature across a range of antibiotic concentrations spanning the full dose-response curve (0, 2, 5, 10, 15, 20µg/mL). The selection rate constant for any temperature by kanamycin concentration, s*rc(t,k)*, was calculated as:

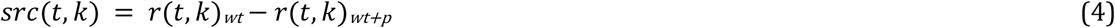

where *r(t,k)_wt_* is the growth rate of the wild-type strain and *r(t,k)_wt+p_* is the growth rate of the wild-type carrying the plasmid (either R1 or RP4). Selection rate constant of 0 indicates no fitness difference between strains. Positive values indicate higher fitness of the wild type, negative values indicate higher fitness of the resistant (plasmid carrying) strain. The selection rate constant calculates absolute differences in growth rate, and allows us to understand how absolute selection is varying across the whole temperature range.

We also estimated the selection coefficient, which is a relative measure of selection that normalises for differences in growth rate driven by temperature, allowing us to isolate the relative impact of kanamycin. The selection coefficient across combinations of temperature and kanamycin concentration, *s(t,k)* was calculated as:

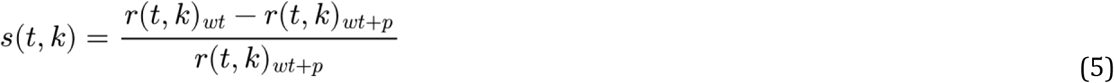

When the coefficient is 0, it means there is no selection for the plasmid. When values are negative, it means that the plasmid is selected for, and positive values indicate the plasmid is selected against (no spread).

We fitted a separate model to both the selection rate constant and selection coefficient as we did for the trait estimates above, but we used a gaussian error structure for the selection coefficient. The response of the model was the estimate of selection and the predictors were temperature (included as a second order polynomial), kanamycin concentration (as a discrete variable), and plasmid, as well as 2-way and 3-way interactions. To demonstrate how selection changed across temperatures, for each kanamycin concentration we calculated the range of temperatures where credible intervals of the predictions do not overlap 0, with values below 0 indicating selection for kanamycin resistance.

#### Analysing invasion assays

To analyse whether our dose response curves can predict outcomes in mixed populations, we carried out invasion assays. We modeled whether the proportion of resistant bacteria (our response) changes across our predictors (all of which are discrete): the two different plasmids (R1 and RP4) temperatures (21°C, 31°C, and 40°C) and kanamycin concentrations (0, 2, 3.5, 5, and 10µg/mL). We used a beta regression and had a random effect of replicate to account for the six different runs of the experiment, and allowed the precision parameter phi to vary with the same factors. To understand when selection occurred, we compared the marginal predictions of the difference in resistance proportion when there is no antibiotic to that of each antibiotic concentration at each temperature. When the 95% credible intervals are greater than 0, then there is selection for resistance compared to no antibiotic.

To analyse how much conjugation occurred in our assays, we modelled the number of transconjugants (CFU per 10 µL) across temperatures and kanamycin concentrations (our predictors as above). We used a negative binomial model with replicate as a random effect, and allowed the dispersion parameter to vary with the same factors. The posterior predictive check of this model demonstrated that it predicted some transconjugant counts that were 2 orders of magnitude higher than the maximum observed, but it was the best model as selected by leave-one-out (loo) comparison against a model where the dispersion parameter varied with (a) just replicate or (b) the same factors but non-interacting. We modelled CFU per 10 µL to keep the counts as close to the observed counts on the plates as possible, which helped with model fitting compared to scaling counts to CFU per mL. The code for fitting these models is in *invasion.R*. We report the median estimate in change of the modeled proportion of resistant bacteria at each kanamycin concentration relative to the drug-free environment using ‘*marginaleffects::contrast()’*.

## Results

### Conjugation rate does not peak at optimal growth temperatures

Bacterial growth and conjugation both had a unimodal response to temperature, however, the thermal performance of conjugation did not follow that of bacterial growth. The drug-free growth rate of the wild-type *E. coli* MG1655 increased to an optimum, *t_opt_*, of 40.5 °C (95% credible intervals = 39.8-41.3) before declining, although growth at the maximum temperature measured (45°C) was still positive and similar to those at 30 °C (Figure 1a).

**Figure 1.**
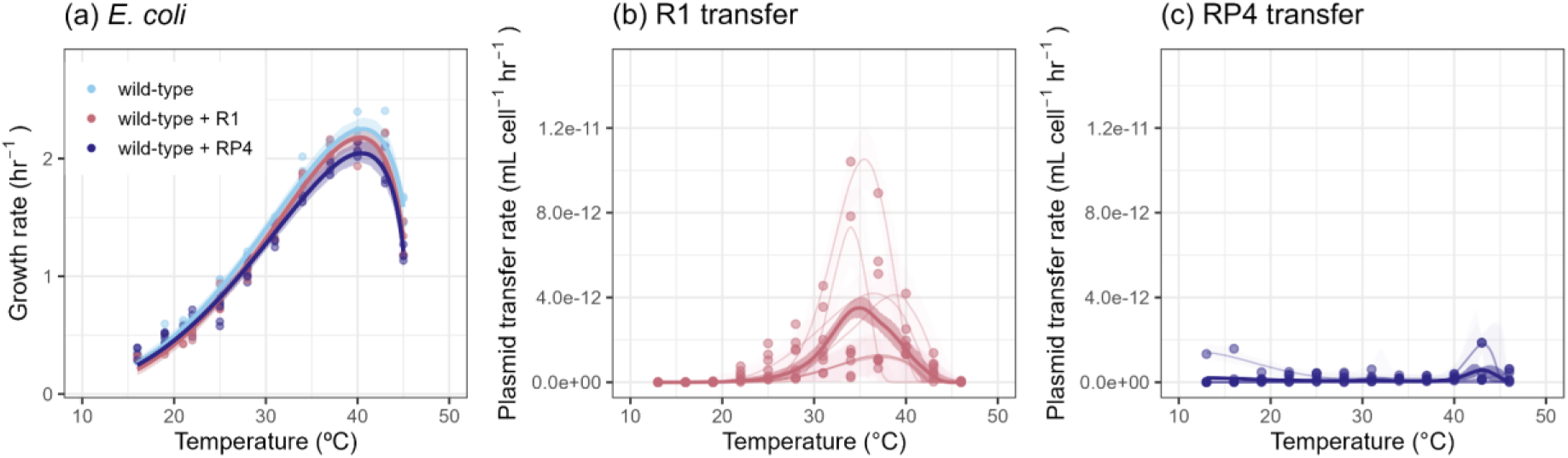
Thermal performance of bacterial growth and plasmid transfer in the absence of kanamycin. (a) Growth of E. coli MG1655 wild-type (light blue) with and without the plasmids R1 (pink) and RP4 (dark blue) increases up to 40 °C before declining. (b) Transfer of the R1 plasmid peaks at 35 °C, but has large variability between replicates. (c) Transfer of the RP4 plasmid is very low across all temperatures. Points represent replicates at each temperature. Solid lines represent the median prediction and shaded bands represent 95% credible intervals. In (b) and (c) darker lines represent population level predictions, and lighter lines represent predictions for each replicate.

The thermal performance of the R1 plasmid transfer was significantly different from the thermal performance of bacterial growth (Figure 1), with the *t_opt_* being 5 °C lower, peaking at 35 °C (34.0-35.9)(Figure 1b). Moreover, R1 plasmid transfer had a higher critical thermal minima (15.8 °C (10.6-21.1)) and lower critical thermal maxima (42.7 °C (40.4-45.7)), resulting in a narrower thermal tolerance of 26.9 °C (20.5-33.2) compared to 46 °C (40.8-51.2) for bacterial growth. These thermal mismatches show that highest plasmid transfer did not occur where growth rates of donor and recipient are maximised, and plasmid transfer only occurs in a restricted section of the bacteria’s thermal tolerance. In contrast to R1, RP4 does not conjugate well in liquid media (48), and we observed very little transfer at any temperatures (Figure 1c). Consequently the model did not converge and we did not obtain reliable parameter estimates.

### Kanamycin impact is lowest at intermediate temperatures

Kanamycin impacted growth of susceptible bacteria (Figure 2), demonstrating a continual curved decline rather than an obvious threshold. We calculated the NSEC, EC_50_, MIC, and size of the sub-MIC selection window for the susceptible bacteria at each temperature (Figure 2 & 3a-d) to quantify how temperature changed the shape of the dose response curve and to test our prediction that susceptibility was lowest at optimum growth temperature.

**Figure 2.**
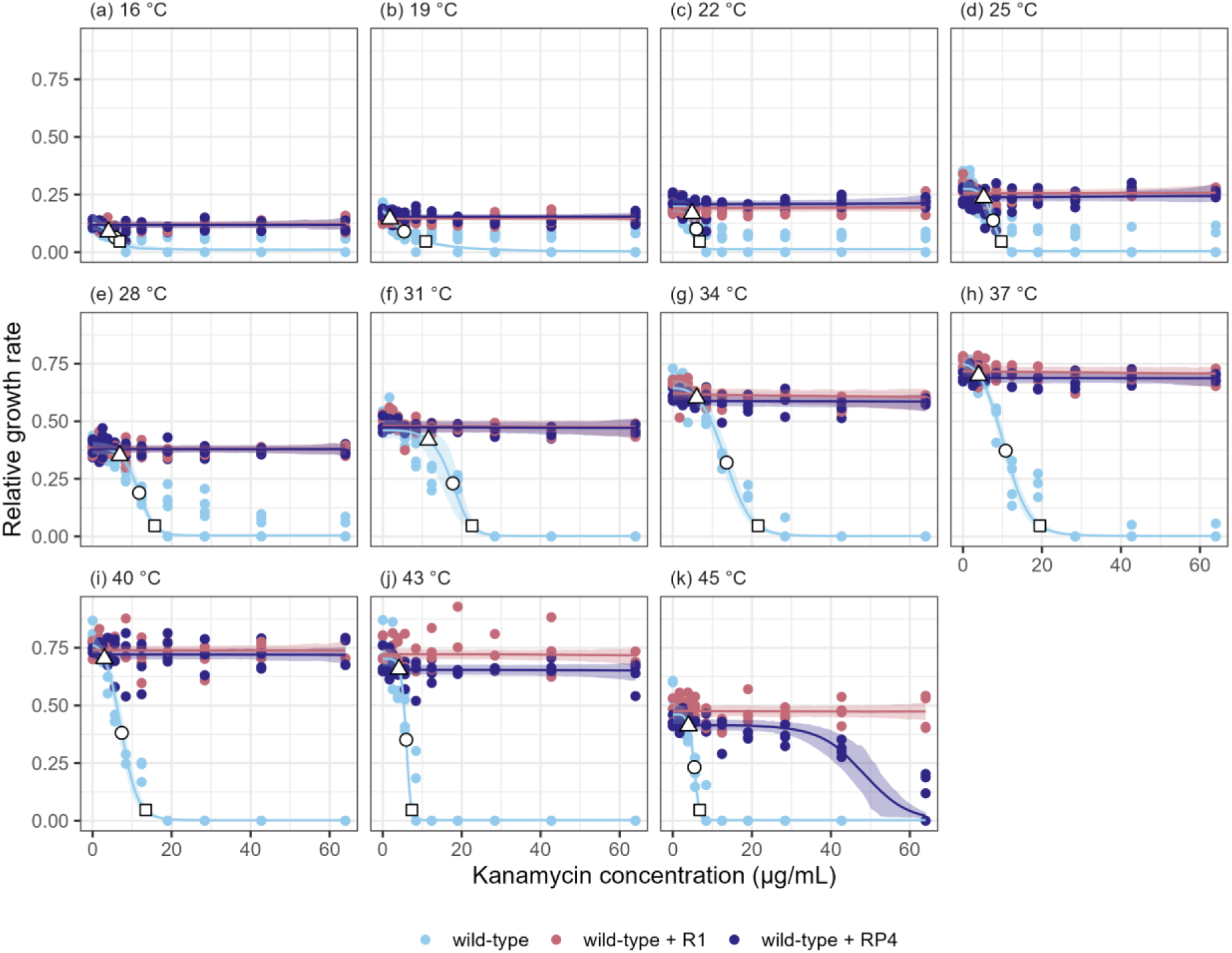
Kanamycin dose response of *E. coli* across temperatures. The relationship between relative growth rate and kanamycin concentration across temperatures for *E. coli* MG1655 without any plasmid (wild-type; light blue), and the same strain containing the plasmids R1 (pink) and RP4 (dark blue). The white points represent key selection traits: NSEC (triangle), EC50 (circle) and MIC (square). Coloured points represent replicates at each temperature. Solid lines represent the median prediction and shaded bands represent 95% credible intervals.

Susceptibility - represented by MIC - decreased with drug-free growth rate up to 31°C, before increasing as temperatures rose towards and beyond *t_opt_* (Figure 3e). The NSEC and MIC of kanamycin for wild-type *E. coli* showed a unimodal response to temperature (Figure 3a,c), but susceptibility was lowest at temperatures well below the optimum temperature for growth (∼40°C), with the highest values for NSEC and MIC at 31.5°C (21.6-45.0°C) and 31.8°C (28.9-35.3°C), respectively. There is considerable uncertainty in these estimates, especially for where NSEC peaks, which includes the maximum temperature assayed. The estimated MIC varied 5-fold (1.50-12.1) across the measured temperature range, ranging from 19 µg/ml (14.2-24.7 µg/mL) to 3.73 µg/ml (1.4-7.9 µg/mL) at 16°C (the lowest measured temperature). A similar fold-change was seen for NSEC (4.03; 1.1-10.8), although absolute changes were smaller. The smallest NSEC of 1.9 µg/mL (0.7-4.3) at 16°C being only 6 µg/mL lower than the highest of 7.7 µg/ml (4.1-13.1). As the MIC changed more in absolute terms than the NSEC, the size of the sub-MIC selection window was also unimodal (Figure 3d), being highest at 32.7°C, with 12.6 µg/mL (8.0-17.5) representing a 5.8 fold change compared to the narrowest window at 16 °C of 2.12 µg/mL (0.7-5.0 µg/mL). Similar quantitative results were found for EC_50_ (Figure 3b).

**Figure 3.**
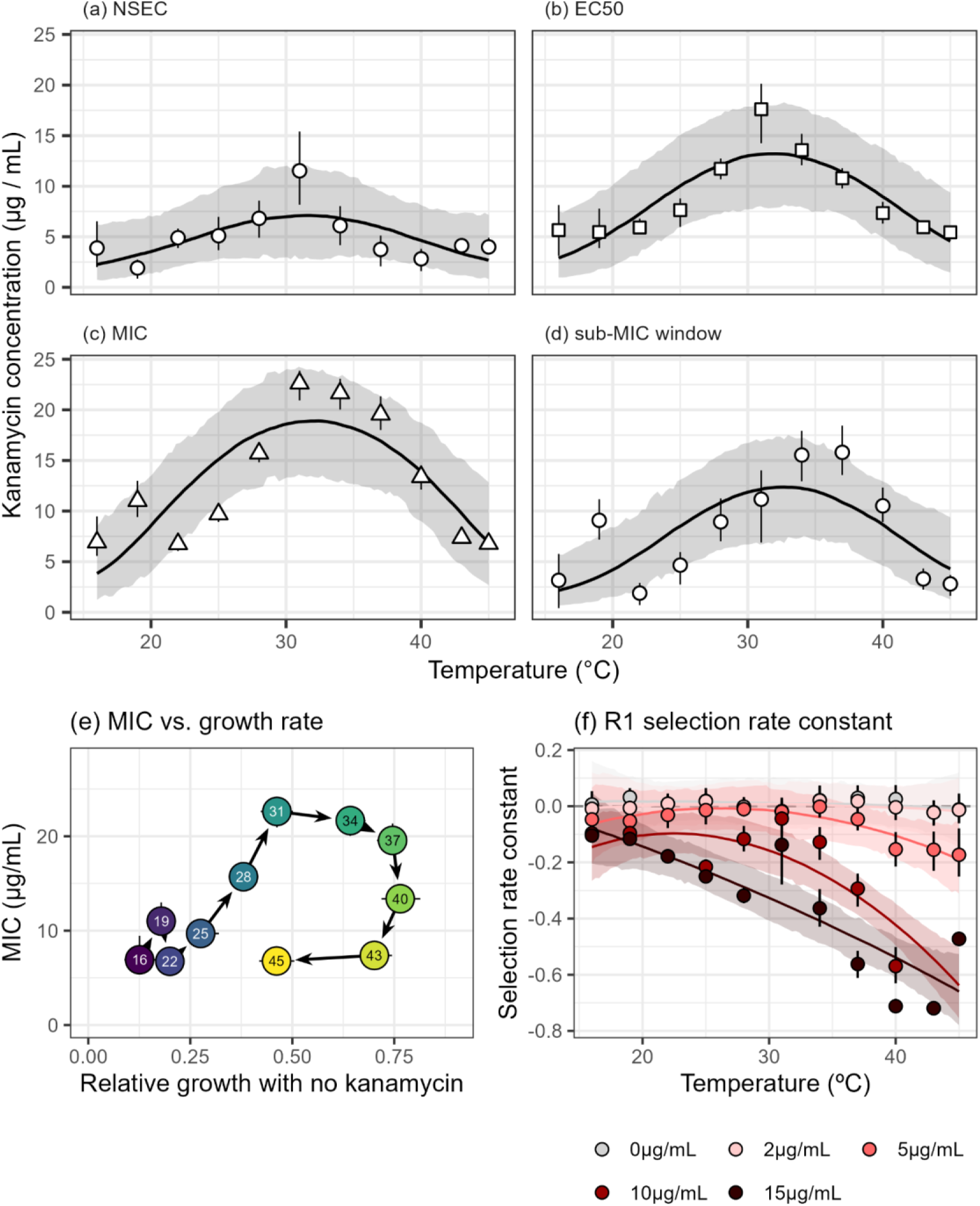
Antibiotic susceptibility and selection changes with temperature. Key antibiotic susceptibility traits (a) NSEC (b) EC50 (c) MIC and (d) the sub-MIC selection window are highest at intermediate temperatures for kanamycin. (e) MIC as a function of kanamycin-free growth rate. (f) Selection increases with temperature, but the relationship between the selection rate constant and temperature changes with kanamycin concentration. In all panels, points and error bars represent median estimates and 95% credible intervals. Lines of best fit and shaded bands show the median and 95% credible intervals of predictions from a model looking at how traits or selection change across temperatures. In (e) arrows link closest temperatures and numbers signify assay temperature.

### Cost of plasmid carriage is negligible

Our experiments allowed us to calculate plasmid cost in multiple ways, all of which estimated no cost to carrying R1, but possibly a weak cost of carrying RP4. First, carrying a plasmid resulted in little change to thermal performance of *E. coli* MG1655, with the *t_opt_* being 40.3 °C (39.6-41.0) for R1, and 40.4 °C (39.7-41.2) for RP4. Carrying RP4 resulted in a lower maximum growth rate (*r_max_*) of 2.05 hr^-1^ (1.96-2.15) compared to 2.25 hr^-1^ (2.16-2.35) and 2.18 hr^-1^ (2.09-2.28) for the wild-type and R1 respectively (Figure 1a). Finally, resistant *E. coli* carrying either R1 or RP4 plasmid had similar growth rates across temperatures and generally did not change with increasing kanamycin concentrations (Figure 2a-i), although wt + RP4 had lower growth rates at 43°C (Figure 2j) and at 46°C bacterial growth was reduced at high kanamycin concentrations (Figure 2k).

### Selection for AMR increases with temperature

Selection for kanamycin resistance increased with warming for the wild-type carrying the R1 plasmid (Figure 3f), with the strongest selection and response to temperature at high kanamycin concentrations. At 15 µg/mL, at or beyond the MIC for susceptible bacteria, selection for resistance occurred at nearly all temperatures, but became stronger with warming. The first temperature at which selection was significant was 17.5°C, where the selection rate constant was -0.1 (-0.2 – -0.001). This increased consistently with warming and was over six times stronger at 45°C, where it was -0.66 (-0.78 – -0.53). At 10 µg/mL, the relationship between temperature and selection was non-linear. Although selection at 45°C was similar to that at 15 µg/mL (-0.64 (-0.76 – -0.50)), and significant selection occurred at 16°C (-0.14 (-0.27 – - 0.006)), at 31°C selection remained similar to those at 16°C (-0.15 (-0.27 – -0.02)). At 5 µg/mL, a concentration between the NSEC and MIC, this same non-linear response resulted in significant selection occurring only at temperatures above 37.4°C. Even at 45°C where selection was strongest, it was only of similar strength to that at cold temperatures for 10 and 15 µg/mL (-0.19 (-0.31 – -0.08)). At concentrations at or near the NSEC (2 µg/mL), we saw no significant selection with all values overlapping 0 (Figure 3f, light red). Similarly, in the absence of antibiotics, credible intervals always overlapped 0, indicating no selection for or against the plasmid (Figure 3f, light grey). The wild-type carrying RP4 showed extremely similar patterns, and this holds for both estimates of selection using selection rate constant and selection coefficient (Figure S15 & S16).

### Selection drives invasion of resistance

Selection for resistance and conjugation rate are both affected by temperature. To explore their relative importance on AMR prevalence, we created mixed populations of plasmid carrying and susceptible bacteria at a range of temperatures and kanamycin concentrations where we expected to see differential selection for AMR and plasmid transfer based on our growth-and plasmid transfer models. If the spread of resistance is primarily driven by selection, we would expect increased prevalence at the hottest temperatures assayed, regardless of plasmid transfer. If horizontal gene transfer was the dominant driver, we would expect elevated AMR prevalence to track temperatures with high plasmid transfer rates.

The proportion of resistance changed across temperatures and kanamycin concentrations (Figure 4a-c), with the largest increase relative to the drug-free environment observed at 40°C despite this temperature having the shortest assay duration, with an 8%, 16% and 68% increase at 3.5, 5 and 15 µg/mL respectively. (Figure 3d-f, Table 1). At 31°C a 44% increase in the proportion of resistant bacteria was observed only at the highest concentration (15µg/mL) where selection was strongest. At 21°C, increases in the proportion of resistant bacteria were evident at 3.5, 5 and 15 µg/mL, with increases of 5%, 3% and 35%. Crucially, the smaller increase in resistant bacteria across kanamycin concentrations at 31°C relative to 40°C occurred despite similar plasmid transfer rates (Figure 1b, Figure 4g-i). This result - alongside observing increases in resistance at 21°C despite nearly no plasmid transfer - strongly suggests that selection is driving these differences (Figure 3f). (Table 1).

**Figure 4.**
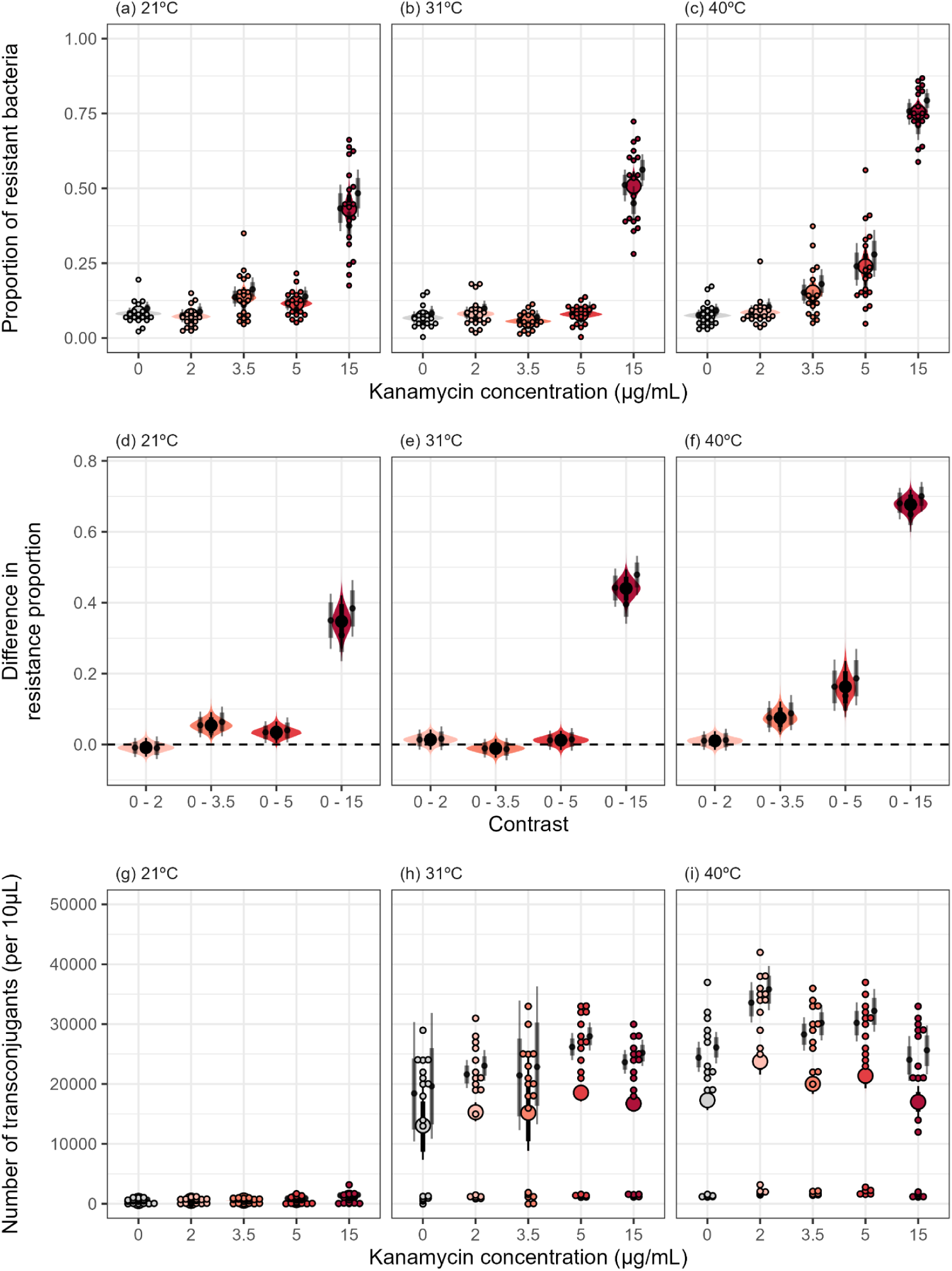
Temperature- and kanamycin-dependent resistance invasion and R1 plasmid transfer in mixed populations. Estimates of the proportion of resistant bacteria across different kanamycin concentrations at (a) 21°C (b) 31°C and (c) 40°C. (d-f) The difference in resistance proportion measured at different kanamycin concentrations compared to zero antibiotics across temperatures. (g-i) Plasmid transfer across temperatures and kanamycin concentrations. Large coloured points and error bars represent marginal median estimates and 95% credible intervals from the models, with shaded violin bands representing the posterior distribution. Small black points and error bars represent median estimates and 95% credible intervals of the replicate level estimates. Small individual points represent individual data points.

**Table 1.** Contrasts of proportion of bacteria carrying R1 and RP4 plasmids at different temperatures and kanamycin concentrations. Contrasts significantly different from 0 are highlighted in bold.

| Plasmid | Temperature (°C) | Contrast (μg/mL) | Median estimate | Lower 95% CI | Upper 95% CI | Different from 0 |
| --- | --- | --- | --- | --- | --- | --- |
| R1 | 21 | 0 - 2 | -0.009 | -0.034 | 0.020 | no |
|  |  | **0 - 3.5** | **0.054** | **0.021** | **0.092** | **yes** |
|  |  | **0 - 5** | **0.034** | **0.006** | **0.066** | **yes** |
|  |  | **0 - 15** | **0.347** | **0.273** | **0.425** | **yes** |
|  | 31 | 0 - 2 | 0.014 | -0.015 | 0.042 | no |
|  |  | 0 - 3.5 | -0.011 | -0.037 | 0.016 | no |
|  |  | 0 - 5 | 0.012 | -0.015 | 0.039 | no |
|  |  | **0 - 15** | **0.440** | **0.387** | **0.492** | **yes** |
|  | 40 | 0 - 2 | 0.011 | -0.015 | 0.037 | no |
|  |  | **0 - 3.5** | **0.075** | **0.037** | **0.123** | **yes** |
|  |  | **0 - 5** | **0.163** | **0.100** | **0.243** | **yes** |
|  |  | **0 - 15** | **0.677** | **0.631** | **0.717** | **yes** |
| RP4 | 21 | 0 - 2 | -0.035 | -0.076 | 0.004 | no |
|  |  | 0 - 3.5 | 0.001 | -0.038 | 0.038 | no |
|  |  | 0 - 5 | 0.027 | -0.019 | 0.076 | no |
|  |  | **0 - 15** | **0.131** | **0.059** | **0.216** | **yes** |
|  | 31 | **0 - 2** | **0.027** | **0.005** | **0.052** | **yes** |
|  |  | 0 - 3.5 | 0.013 | -0.005 | 0.032 | no |
|  |  | **0 - 5** | **0.020** | **0.001** | **0.039** | **yes** |
|  |  | **0 - 15** | **0.255** | **0.194** | **0.328** | **yes** |
|  | 40 | 0 - 2 | 0.004 | -0.024 | 0.036 | no |
|  |  | **0 - 3.5** | **0.031** | **0.000** | **0.066** | **yes** |
|  |  | **0 - 5** | **0.096** | **0.054** | **0.148** | **yes** |
|  |  | **0 - 15** | **0.456** | **0.407** | **0.503** | **yes** |

We saw similar, but less clear cut, patterns in RP4 (Figure S17). At 15µg/mL resistance always invaded, but the proportions were about 20% smaller than for R1 (Table 1). The proportion of resistant bacteria at low concentrations were also smaller than in R1, resulting in weak or no invasion at low antibiotic concentrations across all temperatures (Table 1). Consistent with the conjugation assays, we observed little to no plasmid transfer for RP4 in the invasion assays, reinforcing selection to be the dominant driver of AMR spread under our assay conditions.

## Discussion

We hypothesised that antibiotic selection would be temperature-dependent, emerging from the response of drug-free growth rate to temperature and the negative growth-dependence of susceptibility. We also expected plasmid transfer to increase at higher growth rates. Results from our high-resolution phenotyping showed that selection and spread of kanamycin resistance changed non-linearly with temperature in *E. coli*, but were not always consistent with our above hypotheses. Susceptibility to kanamycin was unimodal with temperature, but was lowest at 31°C, far below optimum growth temperature. Similarly, plasmid transfer peaked at temperatures 5°C below *t_opt_* for growth (Figure 1). We observed stronger selection at higher temperatures, especially at kanamycin concentrations beyond MIC (Figure 3f). In mixed populations of resistant and susceptible bacteria, AMR spread most where selection was strongest (Figure 4), irrespective of plasmid transfer, indicating that selection might be more important for rising AMR at increasing temperatures than HGT.

Temperature changed the shape of the kanamycin dose response (Figure 2). NSEC, MIC, and EC_50_ followed a unimodal pattern, peaking four-to five-fold higher at an intermediate temperature (∼31°C) than at thermal extremes (16°C and 45°C), indicating that the inhibitory effect of kanamycin is weakest at intermediate temperatures (Figure 3a-d). We had expected the weakest effect of kanamycin at *t_opt_*. However, this expectation was based on a study that modulated growth rate via nutrient concentration, a mechanism that likely does not interact with antibiotic efficacy (19). In contrast, *E. coli* was more susceptible to kanamycin above 31°C despite growth rates increasing, consistent with similar physiological interactions seen between hot temperatures and aminoglycosides (18) (Figure 3e). Moreover, when growth rates declined beyond *t_opt_*, kanamycin susceptibility was much higher than lower temperatures with similar growth rates (Figure 3e). This deviation from growth-dependent susceptibility strongly suggests that kanamycin becomes more effective at temperatures above 31°C. This synergistic impact is likely driven by a combination of temperature directly altering antibiotic efficacy, and indirectly through changes in bacterial physiology. Previous work showed heat-shock at 50°C and 55°C combined with exposure to aminoglycosides potentiates antibiotic activity via a combination of increased uptake, irreversible protein aggregation and increased reactive oxygen species (ROS) contributing to aminoglycoside lethality (49). Furthermore, given the overlap in host response to elevated temperature and aminoglycosides, these two stressors may compound one another’s effects on host physiology (18). Regardless of the exact mechanism, temperature shapes the selective landscape for resistance.

Across our growth experiments, we only found a marginal cost of carrying RP4, with no measurable cost of carrying R1. RP4 carrying *E. coli* had a lower *r_max_* compared to the wt, and notably had reduced growth at high kanamycin concentrations when incubated at high temperatures. The mechanism behind this is unknown, but could be due to the increased impact of kanamycin at these temperatures, making the resistance mechanism less effective, or due to high temperatures impacting plasmid stability. Importantly, the metabolic investment during conjugation - a key plasmid cost - cannot be quantified in these assays, but can in our mixed population assays, which we inoculated with 10% plasmid carriers and 90% plasmid free recipients. At the end of the assays, the median proportion of resistance in the absence of antibiotics was 0.07 for R1 and 0.09 for RP4. Not until 5 µg/mL did we see consistent proportions above the 0.1 we inoculated (R1 median = 0.11, RP4 median = 0.147). This suggests a stronger plasmid cost in co-culture than predicted from monoculture, and reinforces the value of labour-intensive but powerful competition experiments and that costs of resistance are difficult to predict across strains and environments (25,50).

Plasmid transfer for the IncFII plasmid R1 peaked at 35°C, 5°C lower than the optimum growth temperature (Figure 1), suggesting factors other than growth rate influence plasmid transfer and its temperature-dependence. Our prediction for plasmid transfer rate was deliberately simplified as many biotic and abiotic factors influence plasmid transfer. Plasmid genetics play a key role in conjugation: R1 and RP4 differ in their transfer regulation (51,52), replication mechanisms and pilus types resulting in large phenotypic differences in our liquid mating assays. This is demonstrated by the low to no transfer observed for the IncP plasmid RP4 across all temperatures (Figure 1c, Figure S17). Conjugation is an energy-intensive process with trade-offs between plasmid transfer that requires expression of transfer machinery and DNA processing, diverting resources from growth (53,54). A temperature that maximises growth rate does not necessarily maximise expression of transfer genes or type IV secretion genes (33,53), pilus assembly and mating pair formation required for plasmid transfer. In general, our work is consistent with plasmid transfer arising from a complex interplay between plasmid-host genetics and ecology, making the response of plasmid transfer to warming difficult to predict (55).

The narrower thermal tolerance of plasmid transfer is not surprising considering that transfer is a complex tightly coordinated process with many potential points of failure (donor regulation, cell contact), resulting in no detectable transfer, while population-growth rate is more predictable under controlled conditions. The narrower thermal tolerance observed for plasmid transfer may also reflect its higher detection limit, since it was measured by plating rather than the more sensitive OD-based growth rate method, and the shorter time frame of the plasmid transfer assays. We tested the relative importance of selection and plasmid transfer for spread of resistance in mixed populations across combinations of temperature and kanamycin concentrations. Overall, we found that selection, not plasmid transfer, was driving invasion of resistance through increases in the number of donors relative to susceptible bacteria (Figure 4). This pattern was consistent across both R1 and RP4, in line with RP4 showing no detectable transfer at any assay temperature and R1 transfer being too low to meaningfully affect the dynamics (Figure S16). For R1, resistance spread less at 31°C, despite similar transfer rates to 40°C, and resistance spread at 21°C despite nearly no transfer. Our results might be quantitatively sensitive to the duration of experiments, and the initial proportions of the assay, but we limited assay duration to avoid fixation of resistance or susceptibility, which can happen with even small fitness differences. Despite running the assays at 21°C for 22 hours (7 times longer than at 40°C), we still saw lower resistance proportions, emphasising the role temperature-dependent growth and selection played in driving the spread of resistance.

How general are these results likely to be? As bacteria generally live below their optimum in the wider environment (7), growth rates are likely to increase with climate warming which will - in turn - increase selection of AMR. This provides a general mechanism through which AMR spreads through populations and communities, and can explain the correlational patterns seen in recent work (1–3). However, the size of the impact of temperature on selection is going to be context-dependent. The shape, sign, and strength of the growth- and temperature-dependence of antibiotic susceptibility will alter the temperature-dependence of selection, which is further complicated by the fundamental impact of temperature on antibiotic efficacy. For example, in contrast to kanamycin, susceptibility to β-lactams and tetracycline increases with growth rate (19,56), which would reinforce the impact of increased growth rates and might result in an even steeper increase in selection with temperature. However, tetracycline also acts in a similar way to cold temperatures (18), which would make it more effective at low temperatures where growth rate is limited, which might counteract this increase. Regardless, even if some antibiotics have a consistent and strong decrease in susceptibility with growth rate, this effect would need to be large to fully counteract the universal temperature-dependent changes in growth. This could be especially true for pathogens such as *E. coli* which generally have high optimum temperatures and can thrive in humans, livestock, and the wider environment (57).

The pattern of more intense selection at high temperature will be dependent on other environmental conditions. First, despite kanamycin acting in a similar way to high temperatures, we saw no selection in the absence of kanamycin at any temperature, providing no additional evidence to the idea that temperature can co-select for resistance (6,58). This suggests that warming may only increase selection when antibiotic pollution is present. Second, our assays were done under nutrient replete conditions where growth rates were maximal, which is rarely the case in the natural environment (59). Under nutrient limited growth, thermal performance curves become shallower (60) which might result in susceptibility being the dominating response of selection rather than drug-free growth rate as can be seen from analysing the selection coefficient. Although increased HGT with temperature might increase AMR prevalence in the absence of selection, this did not happen in our assays even though they were designed to facilitate transfer in liquid culture. Of course, this may change across different plasmids and hosts, but we expect the variation in this to be large and unpredictable compared to the more general expected response of selection which will drive increased AMR irrespective of rates of HGT.

In conclusion, our study demonstrates that the selection and spread of kanamycin resistance increases at higher temperatures due to increased growth rates and antibiotic susceptibility as a result of interactions between kanamycin and temperature. These results highlight that the whole dose response curve, rather than MIC and NSEC alone, should be considered when assessing selection. This is because MIC and NSEC values reflect susceptibility rather than selection, meaning that comparisons across temperatures and other environmental or host conditions that modulate drug-free growth rate may fail to capture differences in selection. Future work should investigate how general these patterns are, and understand how it varies across different antibiotic classes, bacterial species and their combinations with an aim to inform predictions for environmental metagenomic studies. Moreover, experimental evolution provides a strong tool to investigate how temperature-dependent changes in selection alter the evolution and cost of resistance. With increasing evidence that AMR may worsen in a warmer world, it is more important than ever to gain a deeper, mechanistic understanding of how temperature alters the selection, spread, and evolution of antibiotic resistance.

## Acknowledgements

D.P. and Z.E. were funded by the Medical Research Foundation Emerging Leaders Prize (MRF-ELP-ICCH-24-105), and D.P. was further funded by a NERC Independent Research Fellowship (NE/W008890/1). T.D. was funded by a Royal Society University Research Fellowship (URF\R1\231740). We thank Andy Mathews for laboratory assistance and support running the laboratory, and Sarah Ashton, Clare Rule, and Jane Rowe for administrative support. This work would not have been possible without tools generously made available by other researchers.

## Data Availability

The raw and processed data and code to recreate all analyses is publicly available on GitHub (https://github.com/padpadpadpad/https://github.com/padpadpadpad/ecoli_temperature_amr) and will be archived on Zenodo upon publication.

## Supplementary Information for

**Figure S1.**
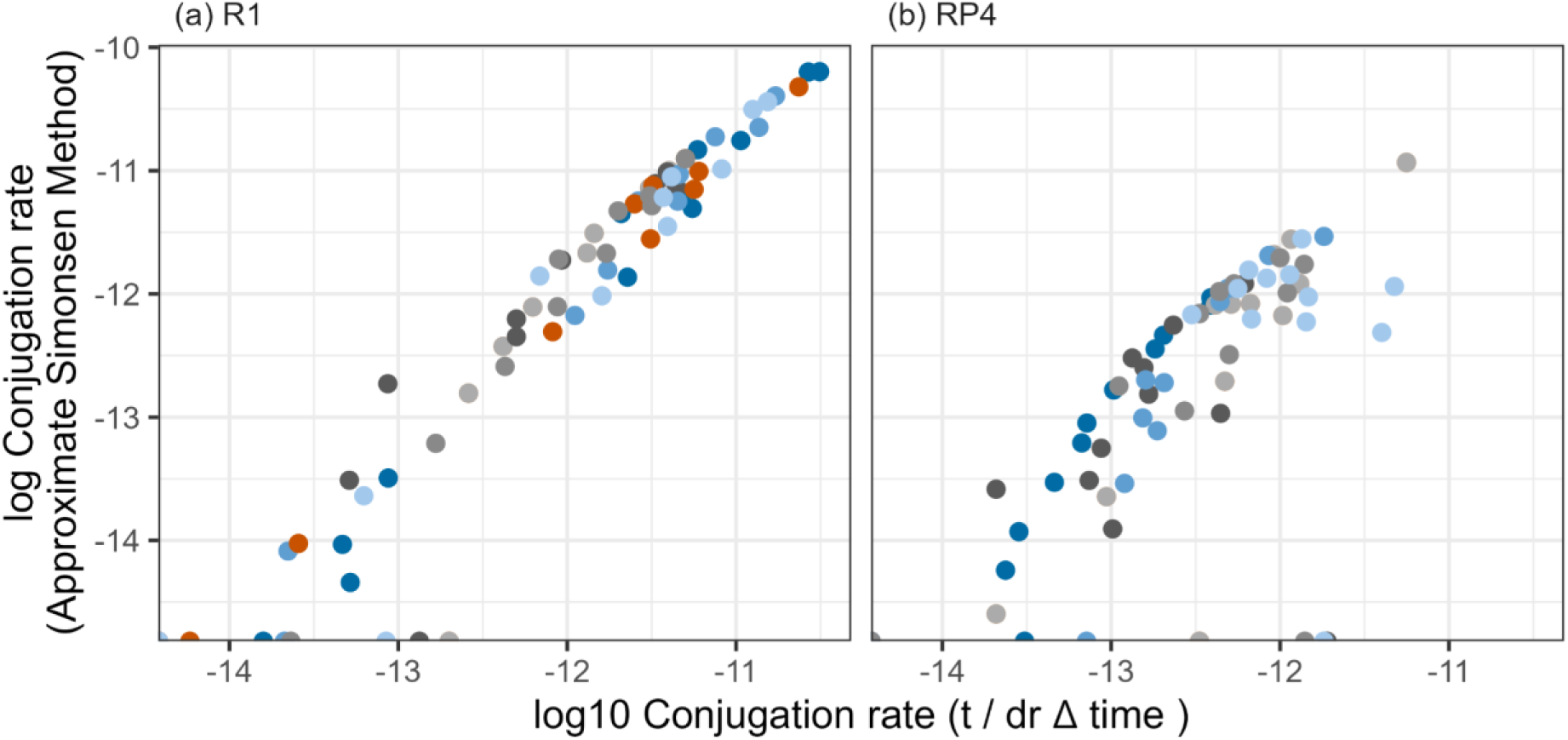
Relationship between different methods of estimating plasmid transfer rate. Plasmid transfer rate was calculated using the gamma method (x axis) and using the Approximate Simonsen Model (y axis) that takes into account growth rates of the donor, recipient, and transconjugants. There is a strong positive relationship between the two methods. Points represent individual estimates. Colours represent different replicates/overnights. Points at the y axis limit represent replicates where the Approximate Simonsen Method failed, which was due to either (a) the gamma estimates also being 0 or (b) poor estimates of the growth rate of donor, recipient, and transconjugant at 46 °C, beyond the range of data collected for the TPCs.

**Figure S2.**
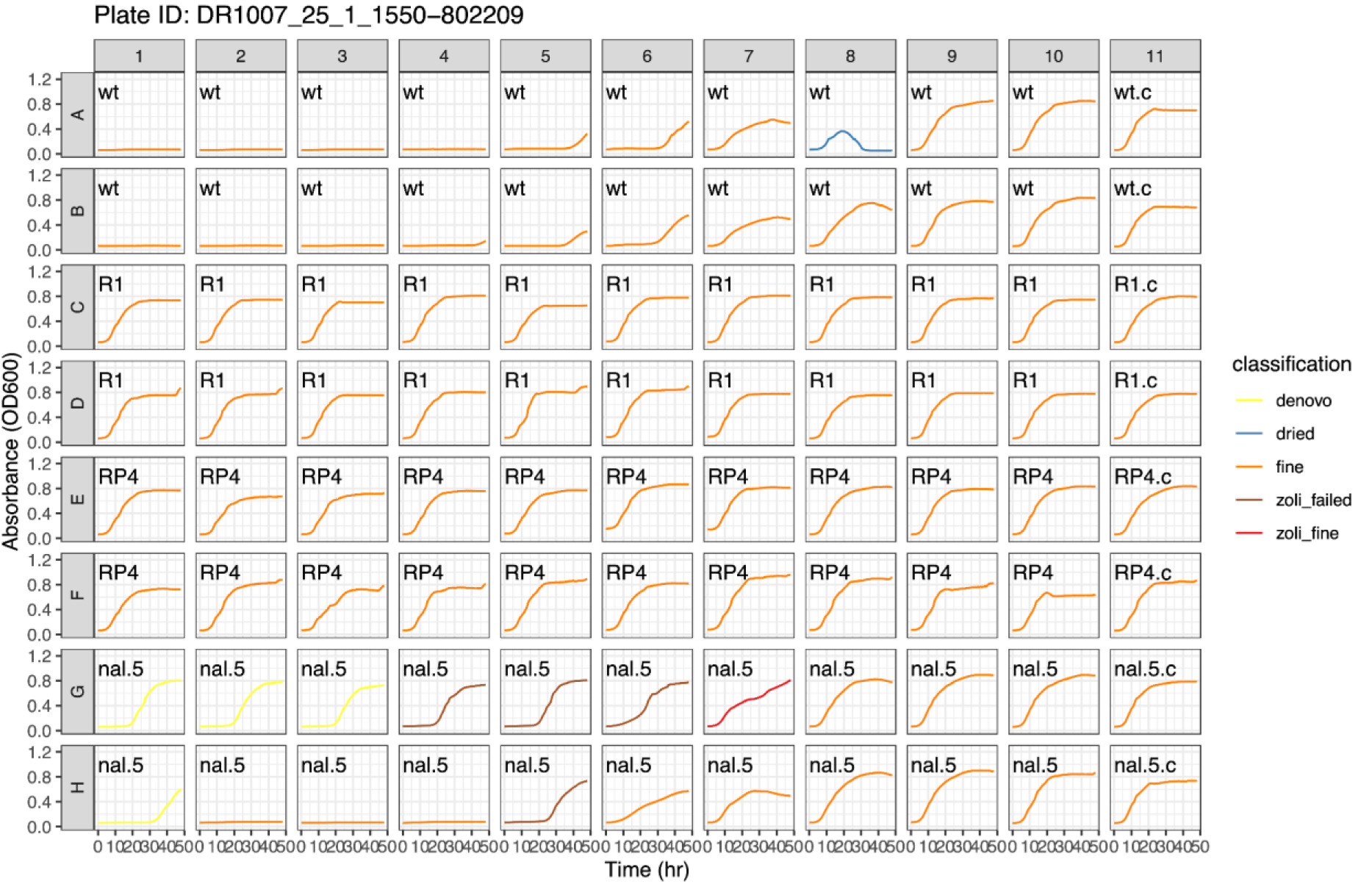
Representative visualisation of cleaning steps for growth rate estimates. Each panel is a well on a 96 well plate. Lines represent growth curves for each well. Colours represent different classifications (see Methods). In brief: fine means the growth rate estimates passed quality control; dried means the well dried up, denovo means we think resistance evolved during the assay, zoli_failed means the well was flagged as failed during the assay, but not programmatically, and zoli_fine the opposite. zoli_fine and zoli_failed are only present where the authors disagreed on outlier assignment. Text inside each panel explains what strain the well contains (wt = wild-type, wt.c = wild-type with no kanamycin, R1 = wild-type + R1 plasmid, RP4 =wild-type + RP4 plasmid, nal.5 = nal^R^ mutant, presence “.c” at the end indicates an absence of kanamycin).

**Table S1.**
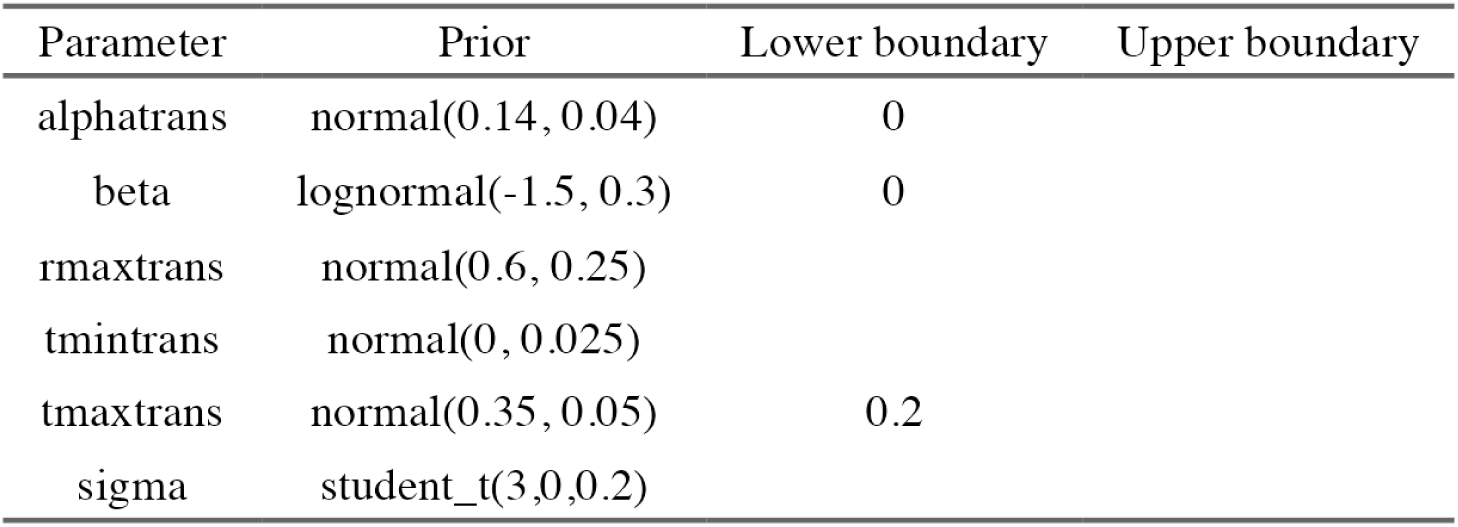
Custom priors for fitting thermal performance curves to bacterial growth data. Priors are reported as they are specified in brms for the transformed model specification. More information on these can be found in *brms_tpc.R*.

**Figure S3.**
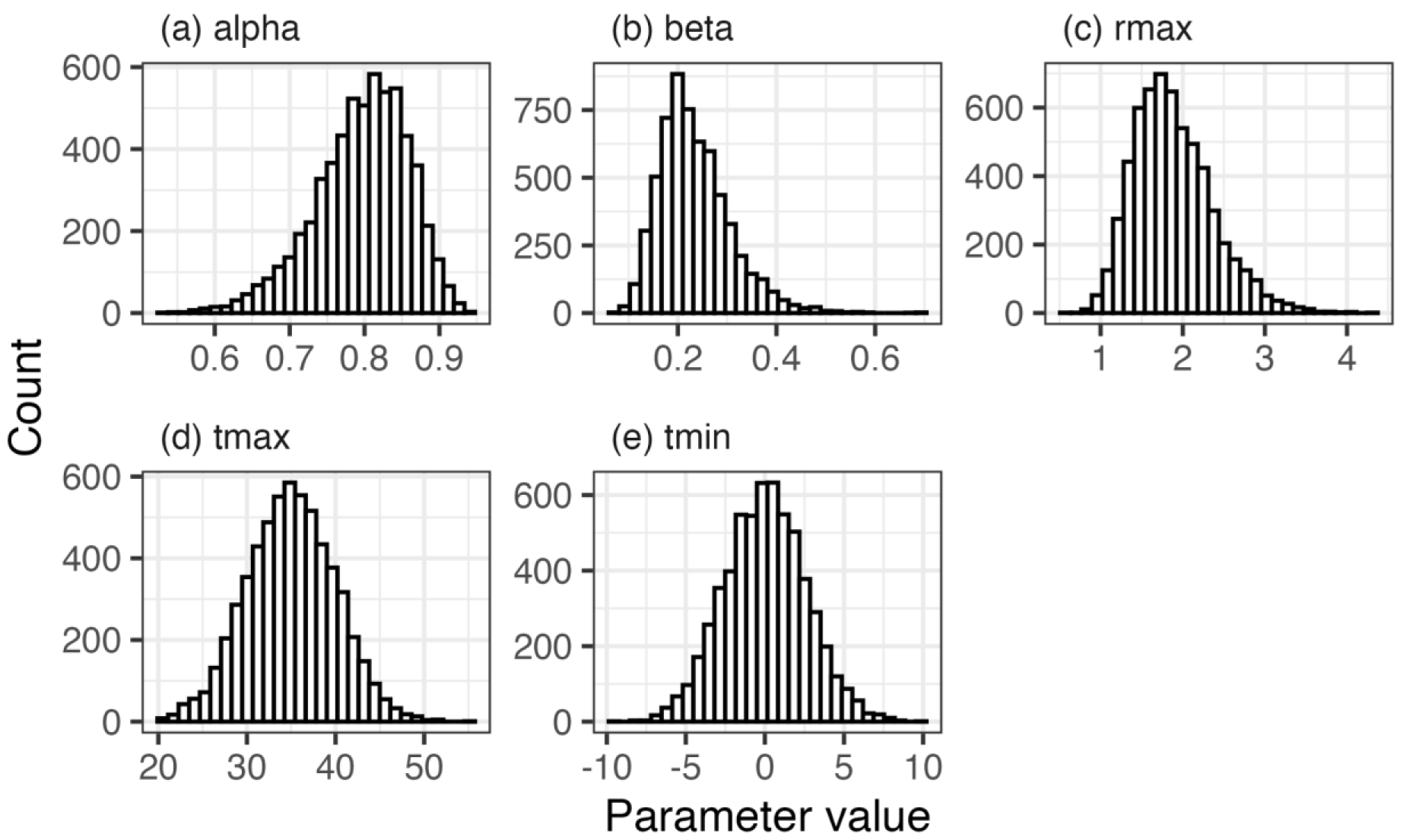
Prior distributions for the thermal performance curves. Prior distributions of back-transformed parameters to show their distribution on the scale of the original flexTPC model formulation. Distributions were created by running the model in the R package ‘brms’ but only sampling from the prior. Only parameters that had custom priors are shown.

**Figure S4.**
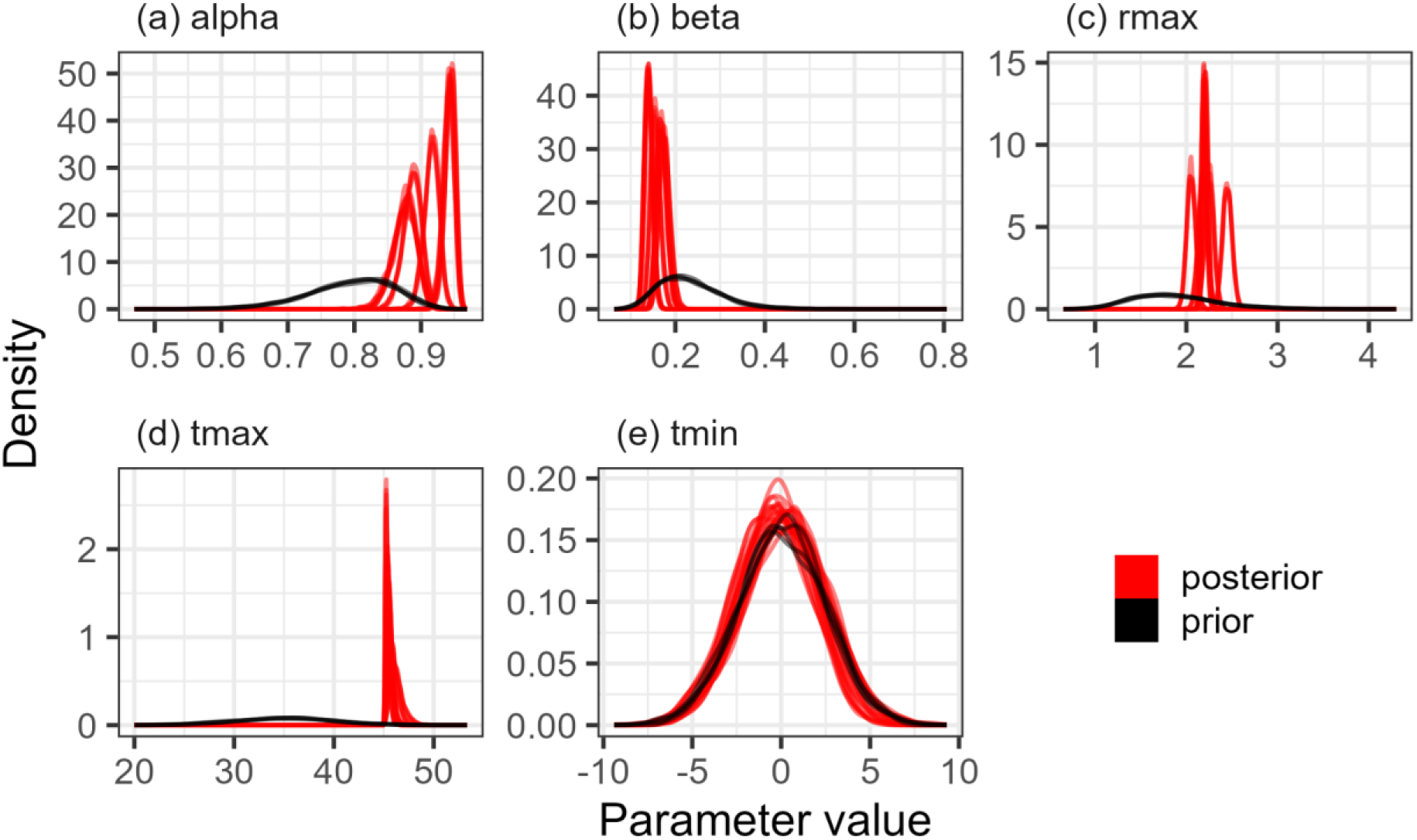
Prior and posterior probability densities for the model of thermal performance of bacterial growth. Probability distributions are backtransformed to their original flexTPC scale. Posterior probability distributions (in red) have moved away from the prior probability distributions (in black), except for tmin where there is very little data. Each line represents an individual chain by strain combination. There are multiple peaks in some parameters because each strain is modelled separately, rather than modelling the deviation from an overall parameter intercept.

**Table S2.**
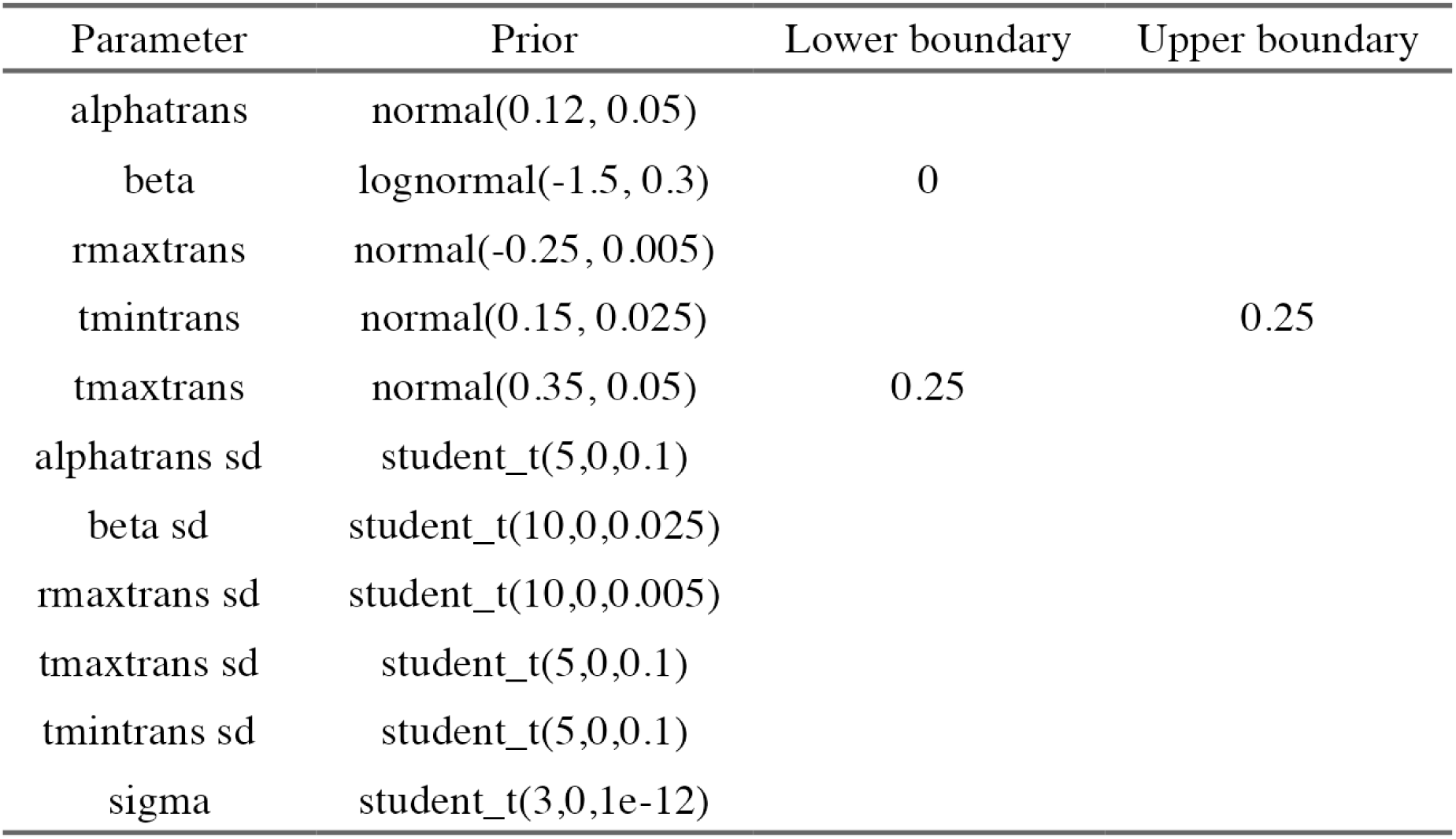
Custom priors for fitting thermal performance curves to plasmid transfer data. Priors are reported as they are specified in brms for the transformed model specification. More information on these can be found in *brms_tpc.R*.

| Parameter | Prior | Lower boundary | Upper boundary |
| --- | --- | --- | --- |
| alphatrans | normal(0.12, 0.05) |  |  |
| beta | lognormal(-1.5, 0.3) | 0 |  |
| rmaxtrans | normal(-0.25, 0.005) |  |  |
| tmintrans | normal(0.15, 0.025) |  | 0.25 |
| tmaxtrans | normal(0.35, 0.05) | 0.25 |  |
| alphatrans sd | student_t(5,0,0.1) |  |  |
| beta sd | student_t(10,0,0.025) |  |  |
| rmaxtrans sd | student_t(10,0,0.005) |  |  |
| tmaxtrans sd | student_t(5,0,0.1) |  |  |
| tmintrans sd | student_t(5,0,0.1) |  |  |
| sigma | student_t(3,0,1e-12) |  |  |

**Figure S5.**
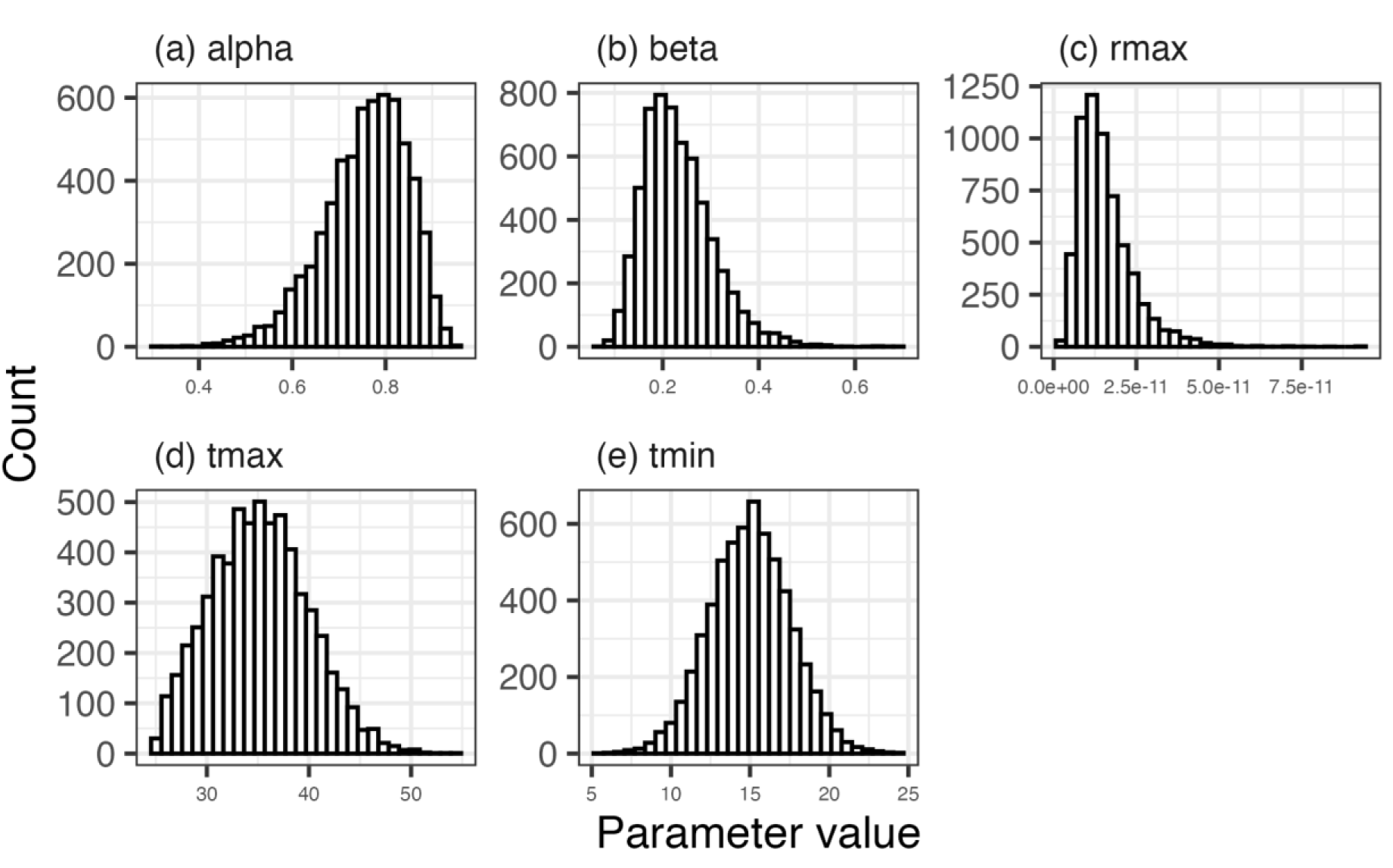
Prior distributions of the fixed effects for the thermal performance of plasmid transfer model. Prior distributions of back-transformed parameters to show their distribution on the scale of the original flexTPC model formulation. Distributions were created by running the model in the R package ‘brms’ but only sampling from the prior. Only parameters that had custom priors are shown.

**Figure S6.**
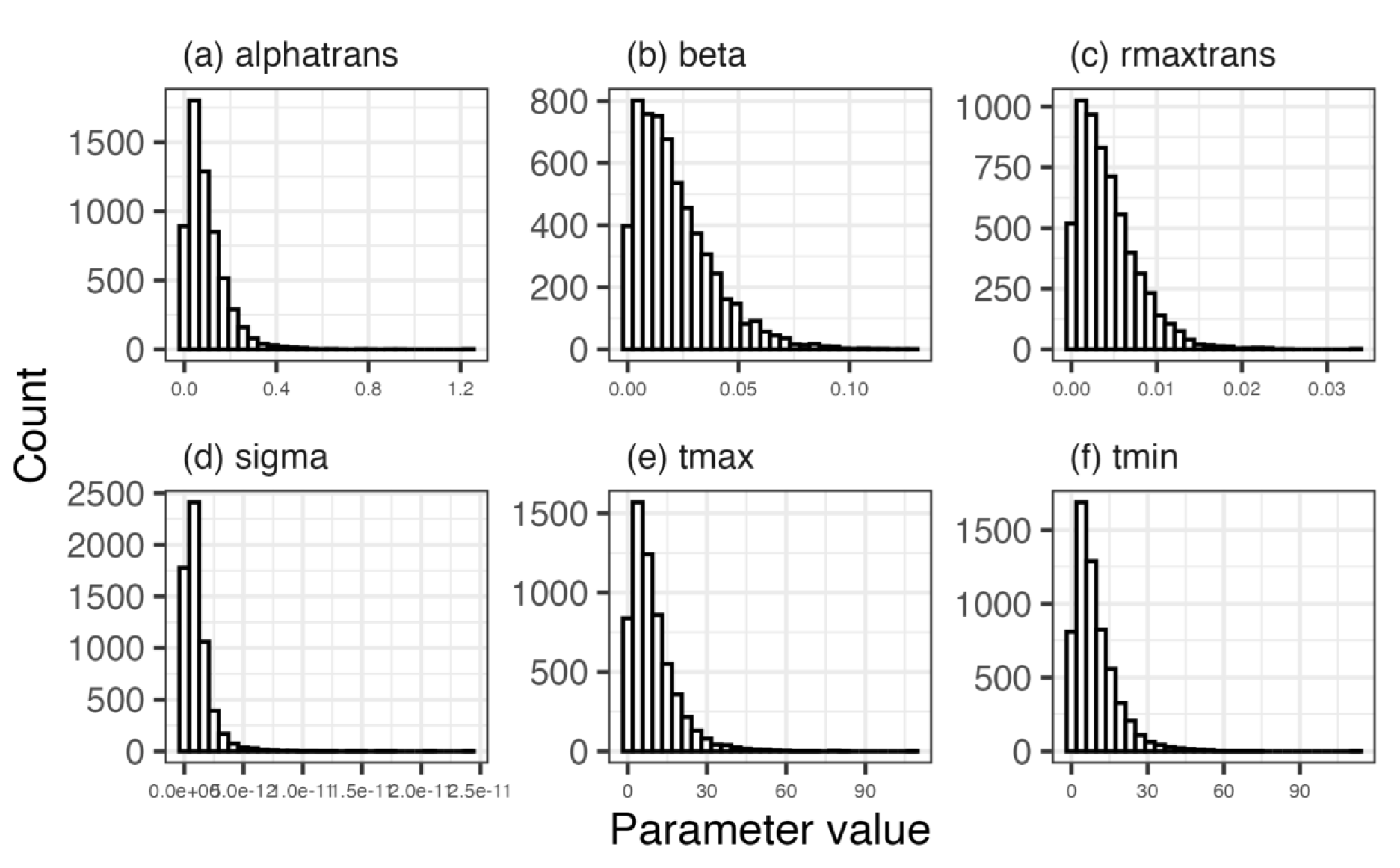
Prior distributions for the random effects (and sigma) of the thermal performance of plasmid transfer model. Where appropriate, prior distributions of back-transformed random effect parameters to show their distribution on the scale of the original flexTPC model formulation. Distributions were created by running the model in the R package ‘brms’ but only sampling from the prior. Only parameters that had custom priors are shown.

**Figure S7.**
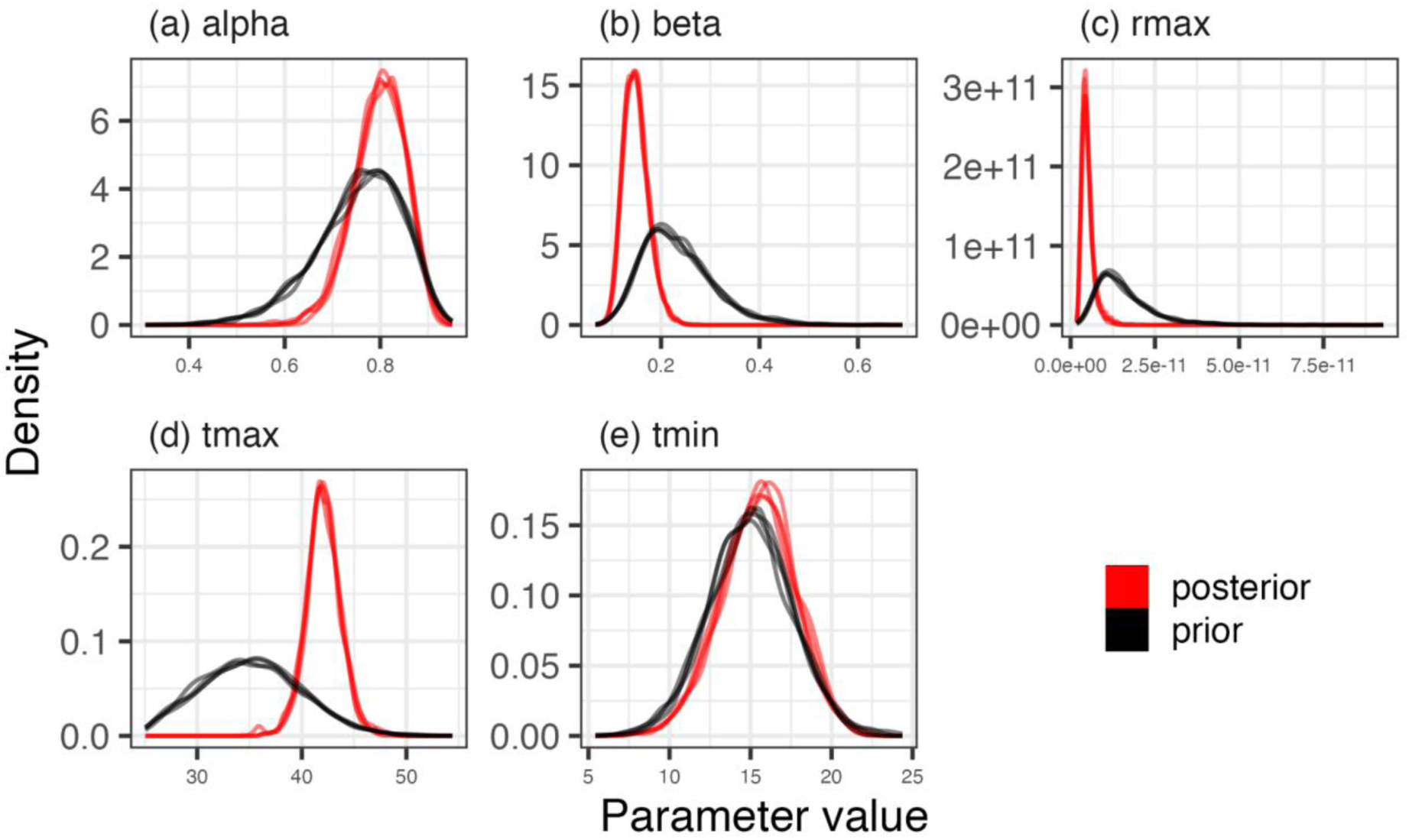
Prior and posterior probability densities for fixed effect parameters of the model of thermal performance of R1 plasmid transfer. Probability distributions are backtransformed to their original flexTPC scale. Posterior probability distributions (in red) have moved away from the prior probability distributions (in black), except for tmin where there is very little data. Each line represents an individual chain.

**Figure S8.**
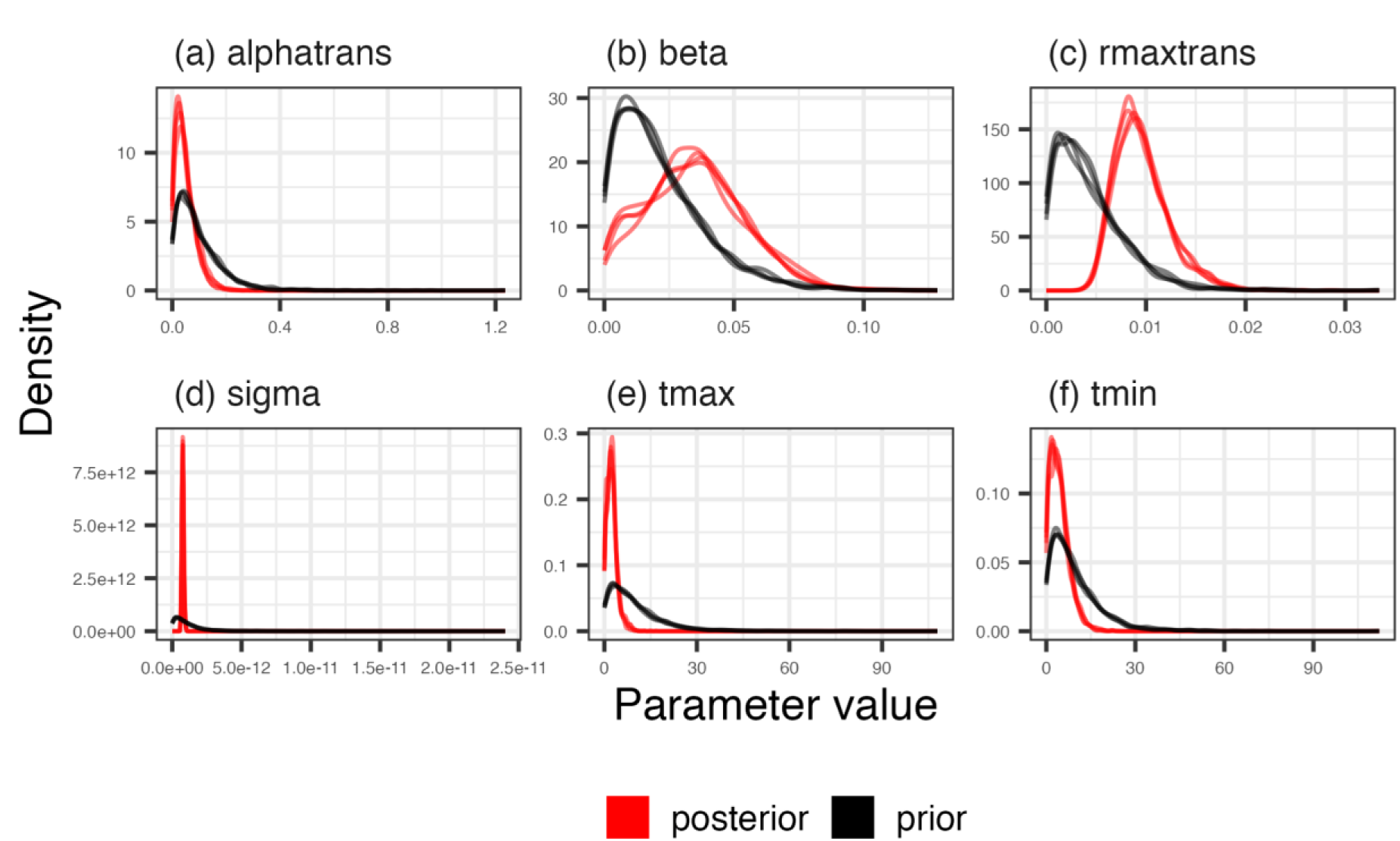
Prior and posterior probability densities for random effect parameters (and sigma) of the model of thermal performance of R1 plasmid transfer. Probability distributions are backtransformed to their original flexTPC scale. Posterior probability distributions (in red) have moved away from the prior probability distributions (in black). Each line represents an individual chain.

**Figure S9.**
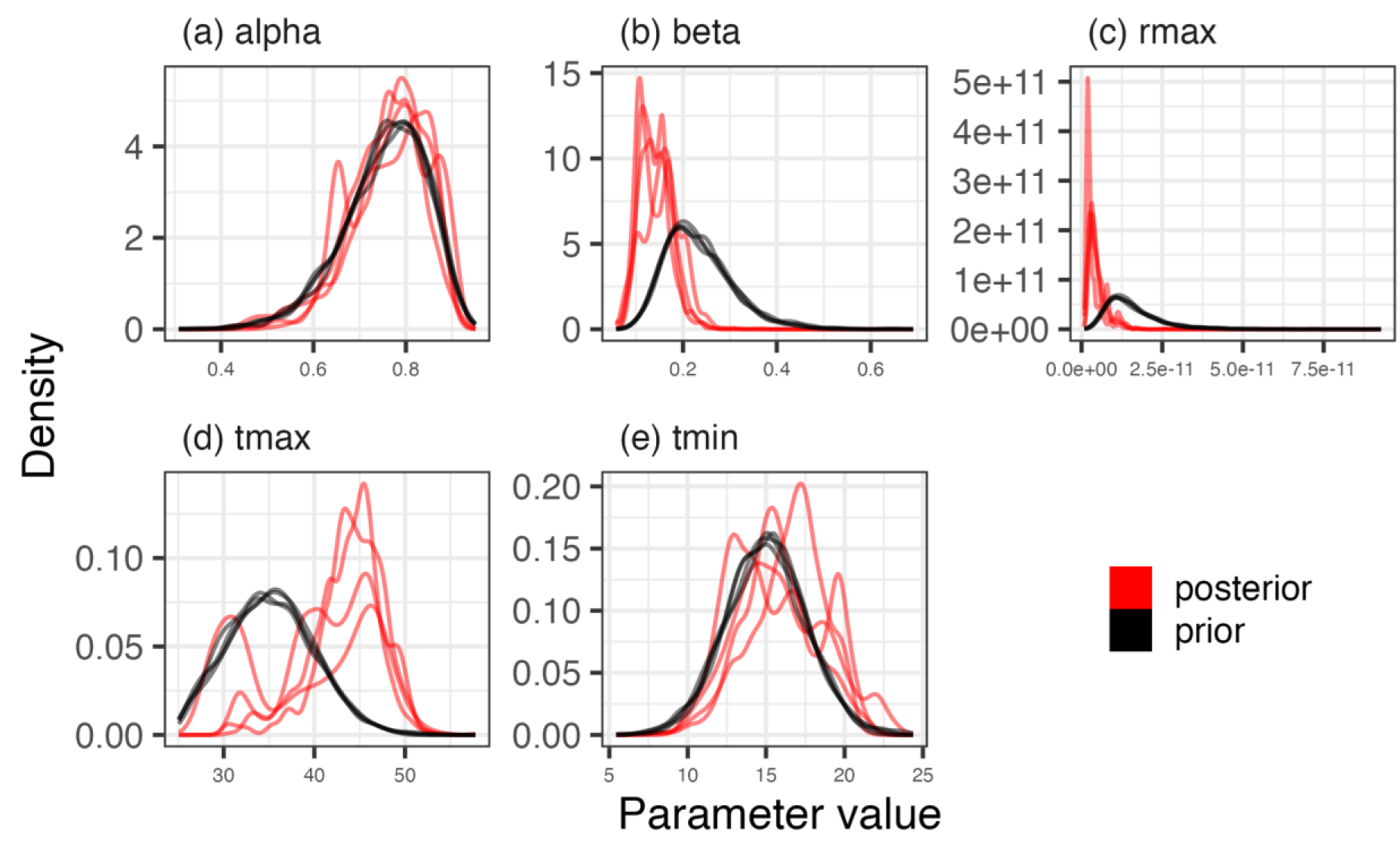
Prior and posterior probability densities for fixed effect parameters of the model of thermal performance of RP4 plasmid transfer. Probability distributions are backtransformed to their original flexTPC scale. Posterior probability distributions (in red) have not converged overly well, and alpha and tmin have not moved much away from the prior probability distributions (in black). Each line represents an individual chain.

**Figure S10.**
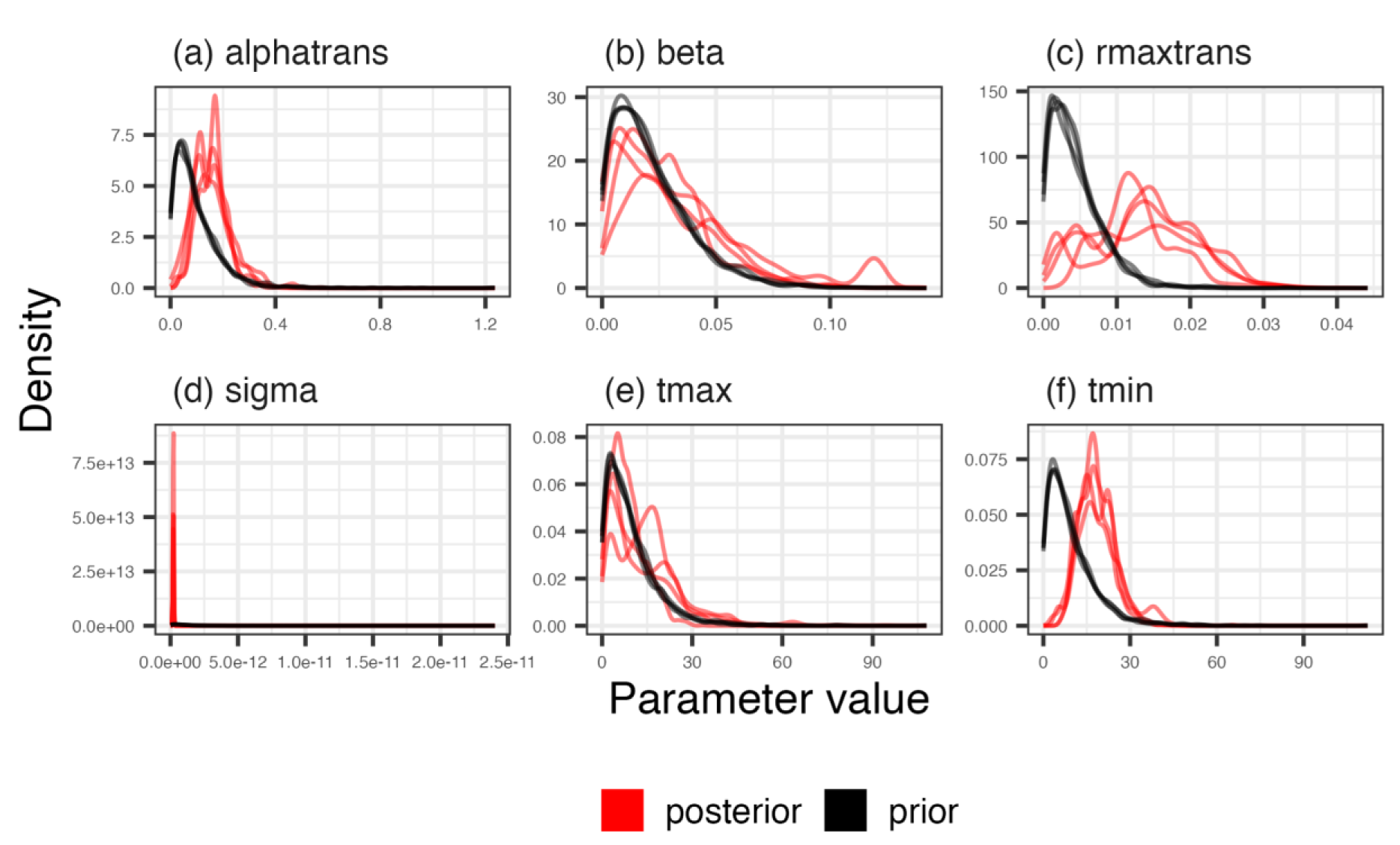
Prior and posterior probability densities for random effect parameters (and sigma) of the model of thermal performance of RP4 plasmid transfer. Probability distributions are backtransformed to their original flexTPC scale. Posterior probability distributions (in red) have not converged overly well, and tmax and beta have not moved much away from the prior probability distributions (in black). Each line represents an individual chain.

**Table S3.** Parameter estimates for the thermal performance curves of E. coli MG1655 without (wt) and with the plasmids R1 (wt + R1) and RP4 (wt + RP4).

| Thermal performance | TPC parameter | Median estimate | Lower 95% CI | Upper 95% CI |
| --- | --- | --- | --- | --- |
| wt | alpha | 0.88 | 0.84 | 0.91 |
|  | beta | 0.18 | 0.15 | 0.2 |
|  | rmax | 2.25 | 2.16 | 2.35 |
|  | thermal tolerance | 46.02 | 40.84 | 51.2 |
|  | tmax | 46.2 | 45.31 | 47.67 |
|  | tmin | 0.26 | -4.55 | 4.87 |
|  | topt | 40.54 | 39.83 | 41.32 |
| wt + R1 | alpha | 0.88 | 0.84 | 0.91 |
|  | beta | 0.17 | 0.15 | 0.19 |
|  | rmax | 2.18 | 2.09 | 2.28 |
|  | thermal tolerance | 45.63 | 40.88 | 50.6 |
|  | tmax | 45.87 | 45.21 | 46.9 |
|  | tmin | 0.33 | -4.37 | 4.75 |
|  | topt | 40.27 | 39.57 | 41.06 |
| wt + RP4 | alpha | 0.89 | 0.86 | 0.91 |
|  | beta | 0.17 | 0.15 | 0.19 |
|  | rmax | 2.05 | 1.96 | 2.15 |
|  | thermal tolerance | 45.83 | 41.24 | 50.47 |
|  | tmax | 45.55 | 45.13 | 46.25 |
|  | tmin | -0.22 | -4.56 | 4.36 |
|  | topt | 40.43 | 39.72 | 41.19 |
| R1 transfer | alpha | 0.78 | 0.67 | 0.87 |
|  | beta | 0.14 | 0.1 | 0.19 |
|  | rmax | 4.1e-12 | 3.5e-12 | 4.6e-12 |
|  | thermal tolerance | 26.94 | 20.45 | 33.21 |
|  | tmax | 42.7 | 40.39 | 45.72 |
|  | tmin | 15.84 | 10.58 | 21.05 |
|  | topt | 34.96 | 34.1 | 35.88 |
| RP4 transfer | alpha | 0.69 | 0.52 | 0.85 |
|  | beta | 0.14 | 0.06 | 0.22 |
|  | rmax | 1.5e-12 | 7.7e-13 | 5.3e-12 |
|  | thermal tolerance | 26.36 | 12.66 | 38.6 |
|  | tmax | 45.6 | 37.48 | 52.93 |
|  | tmin | 19.43 | 9.57 | 27.74 |
|  | topt | 43.09 | 30.39 | 45.27 |

**Table S4.** Custom priors for fitting dose response curves. Priors are reported as they are specified in brms for the transformed model specification. More information on these can be found in *brms_drc.R*.

| Parameter | Prior | Lower boundary | Upper boundary |
| --- | --- | --- | --- |
| betatrans | normal(0, 0.5) |  |  |
| bot | beta(2, 5) | 0 | 1 |
| top | beta(5, 2) | 0 | 1 |
| ec50trans | normal(0.275, 0.1) | 0.001 |  |
| bot sd | student_t(3,0,0.25) |  |  |
| top sd | student_t(3,0,0.25) |  |  |
| ec50trans sd | student_t(5,0,0.25) |  |  |
| betatrans sd | student_t(5,0,0.25) |  |  |
| phi | gamma(0.01, 0.01) |  |  |

**Figure S11.**
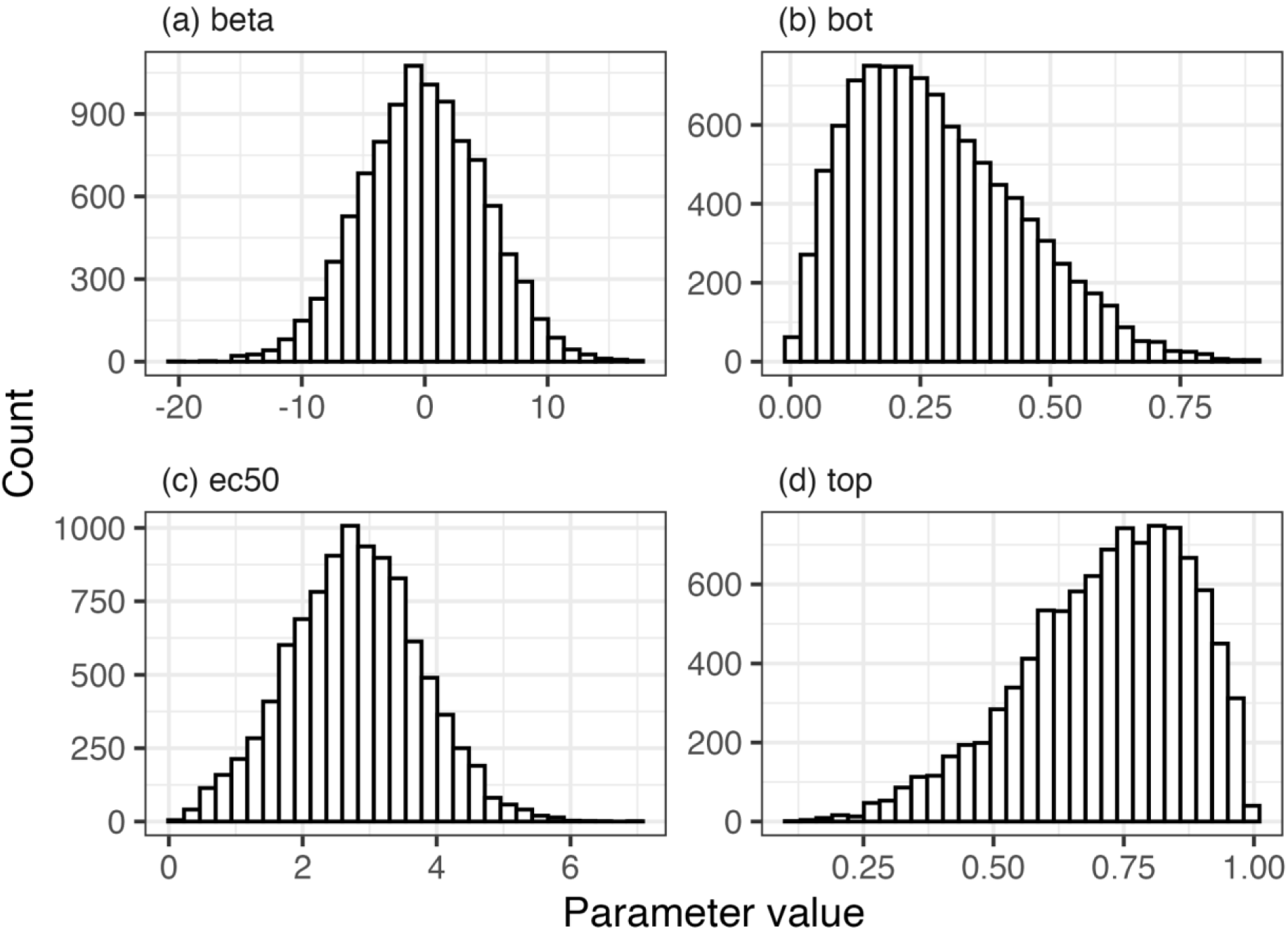
Prior distributions for the fixed effects of the dose response model. Prior distributions are back-transformed to show their distribution on the scale of the original dose response model formulation. Distributions were created by running the model in the R package ‘brms’ but only sampling from the prior.

**Figure S12.**
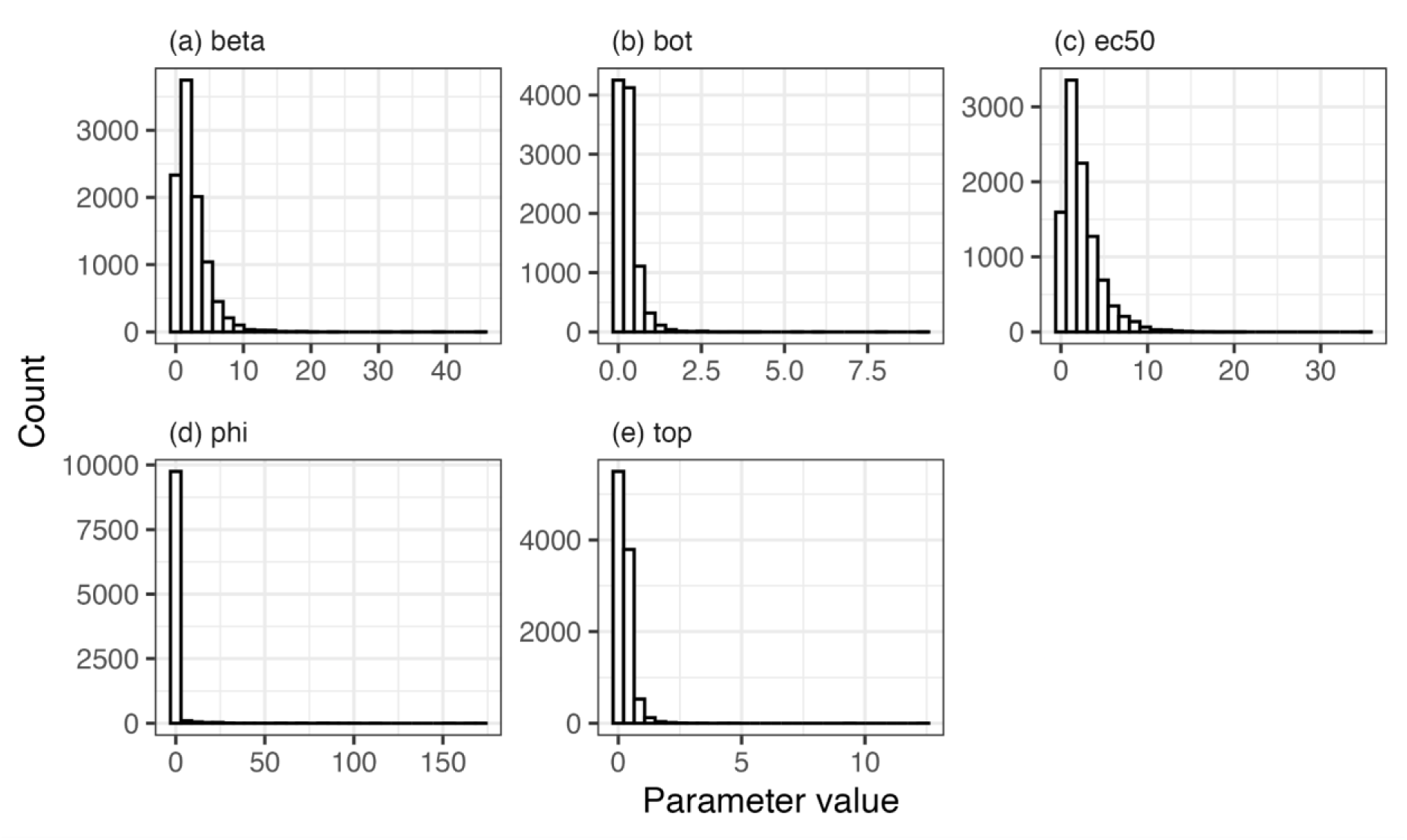
Prior distributions for the random effects of the dose response model. Where needed, prior distributions are back-transformed to show their distribution on the scale of the original dose response model formulation. Distributions were created by running the model in the R package ‘brms’ but only sampling from the prior.

**Figure S13.**
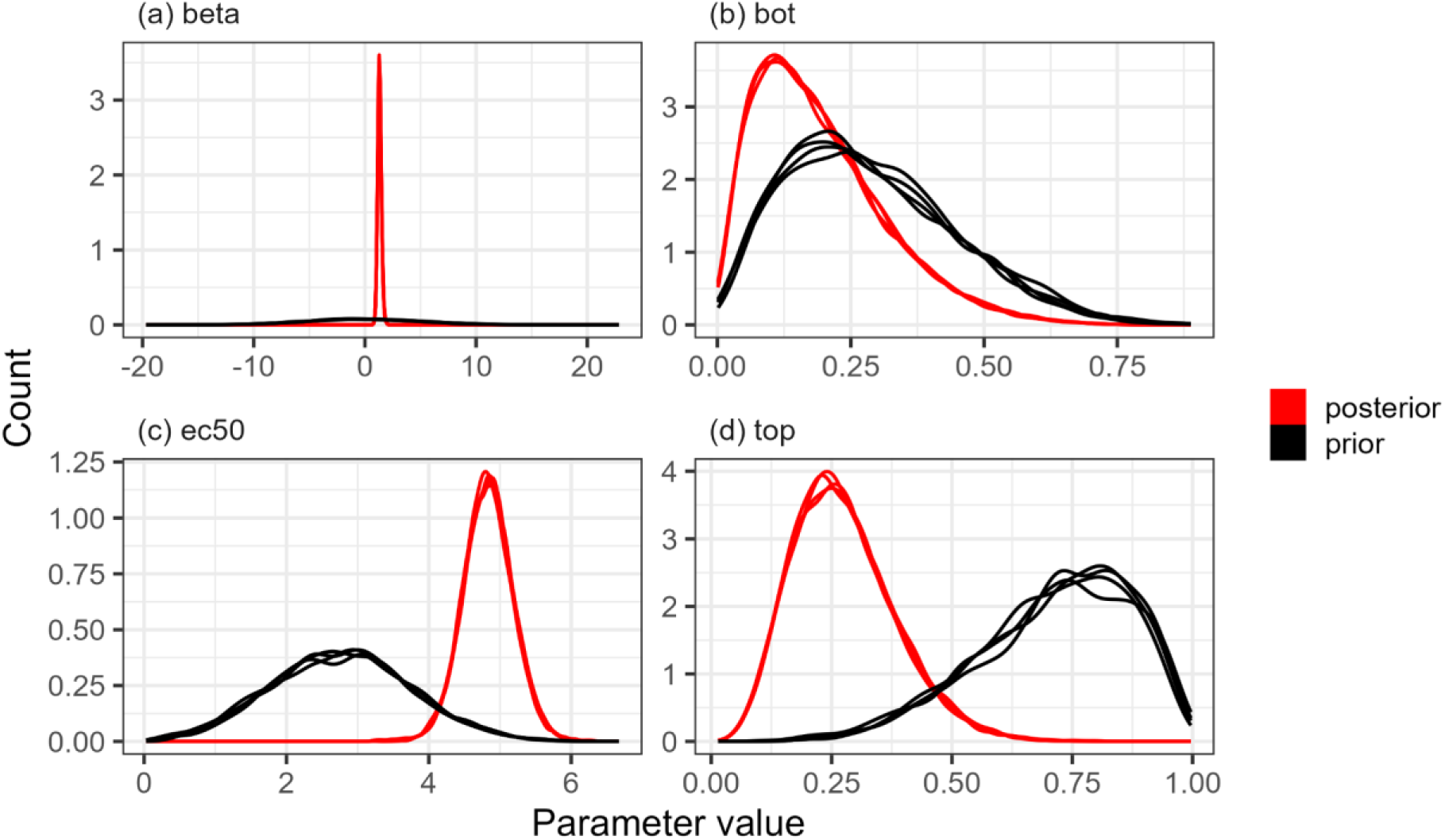
Prior and posterior probability densities for the fixed effect parameters of the dose response curve model. Probability distributions are backtransformed to their original scale. Posterior probability distributions (in red) have moved away from the prior probability distributions (in black). Each line represents an individual chain.

**Figure S14.**
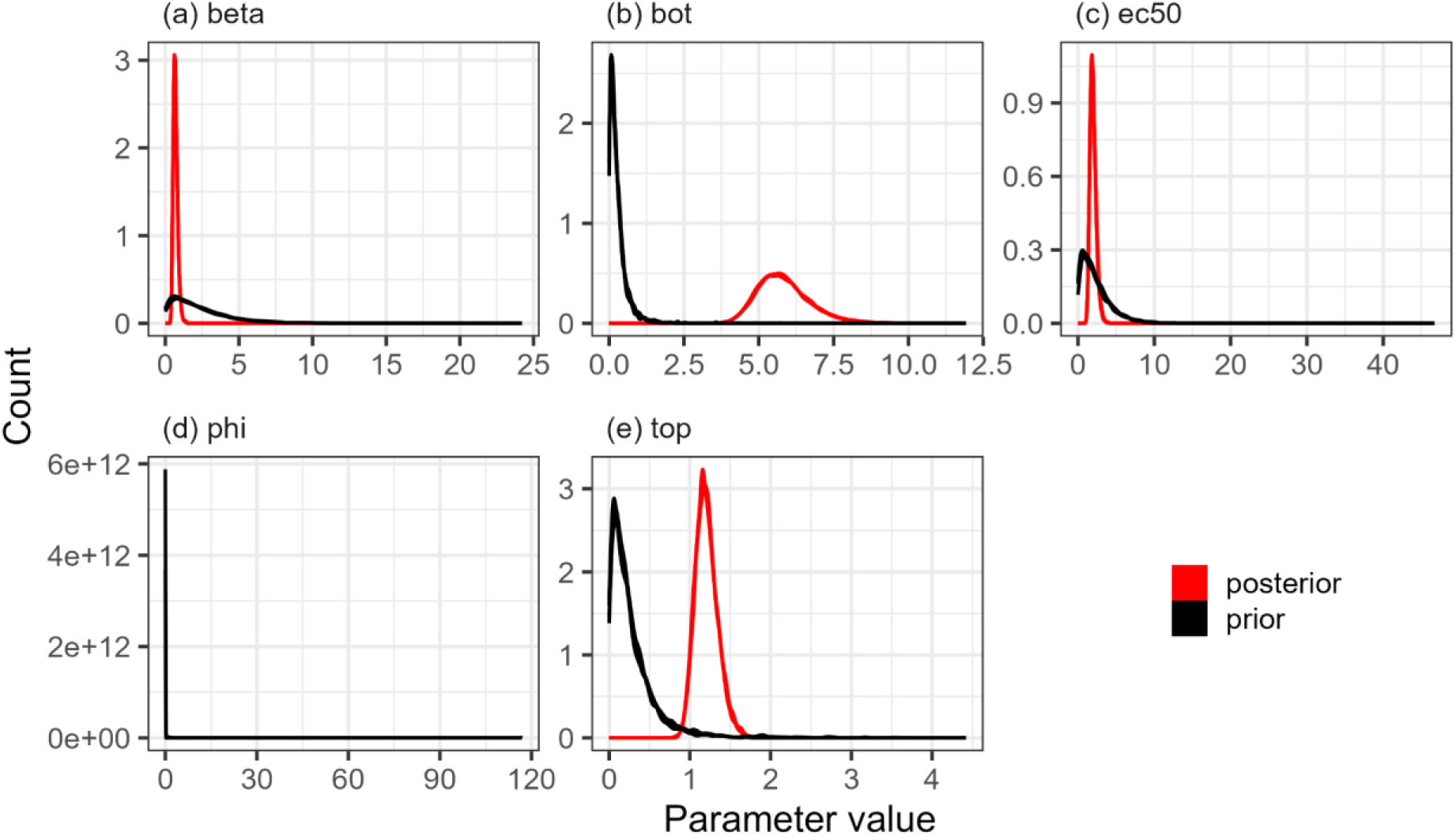
Prior and posterior probability densities for the random effect parameters (and phi) of the dose response curve model. Probability distributions are backtransformed to their original scale. Posterior probability distributions (in red) have moved away from the prior probability distributions (in black). Each line represents an individual chain.

**Table S5.** Custom prior for fitting a polynomial to dose response trait data. Only custom priors are shown. The sigma prior is weakly informative and based on a model fit to the whole dataset without using a truncated normal distribution. More information on these can be found in *brms_drc.R*.

| Parameter | Prior | Lower boundary | Upper boundary |
| --- | --- | --- | --- |
| sigma | normal(3.15, 2) |  |  |

**Table S6.** Custom prior for fitting a beta regression to proportion of resistant *E.coli* and negative binomial to count data of invasion dataset. More information on these can be found in *invasion.R*.

| Parameter | Prior | Lower boundary | Upper boundary |
| --- | --- | --- | --- |
| Intercept | student_t(3, 0, 2.5) |  |  |
| b | normal(0, 2.5) |  |  |

**Figure S15.**
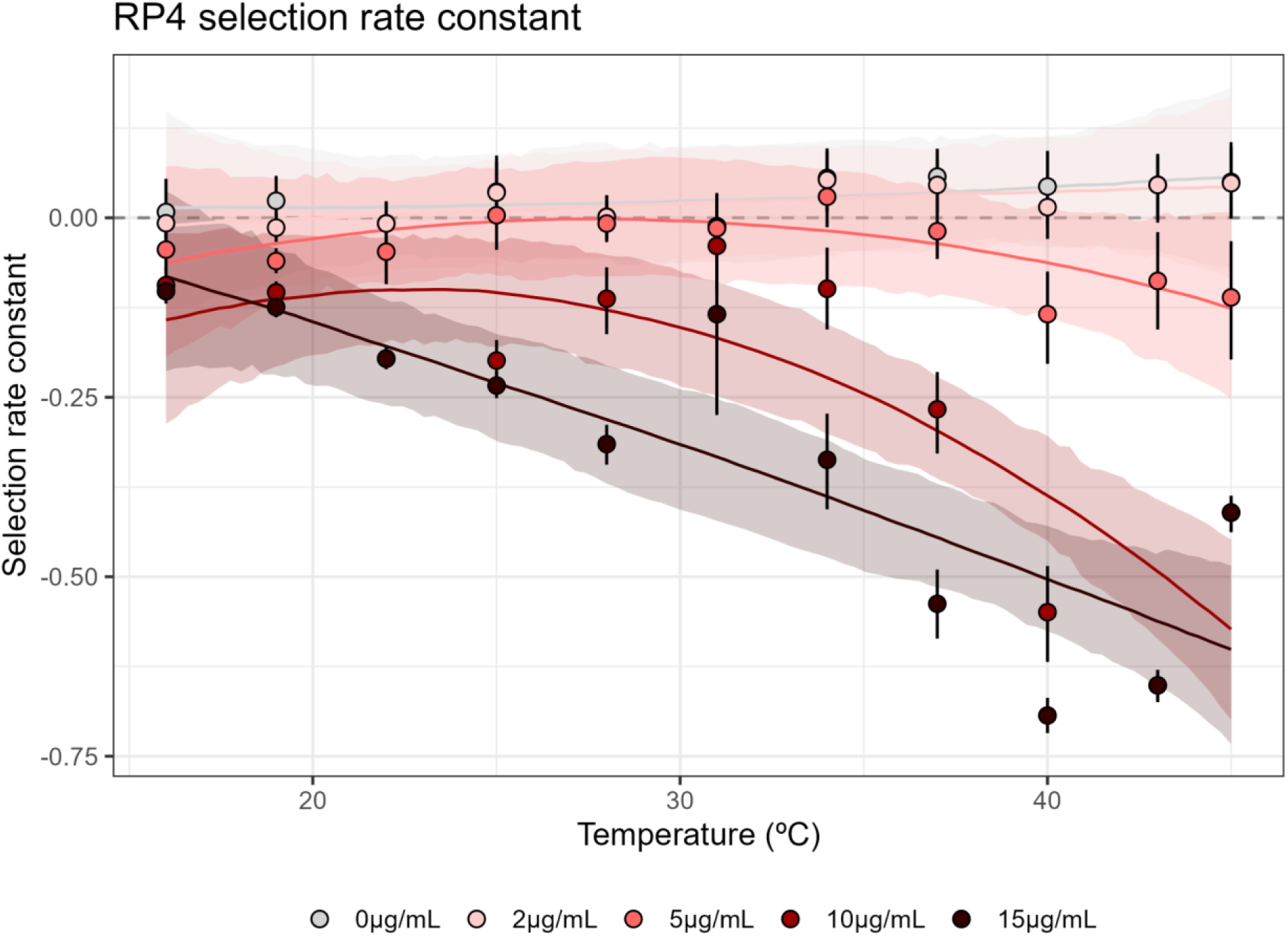
Absolute selection is strongest at high temperatures in RP4. Negative values reflect selection for the plasmid, and positive values reflect selection against. Selection changes non-linearly with changes in antibiotic concentration and temperature. At concentrations at or beyond the MIC (15µg/mL & 10µg/mL), selection increases with temperature. At lower concentrations there is minimal selection at any temperature, but weak selection - if it does occur - happens at the highest and lowest temperatures. Points and error bars represent median estimates and 95% credible intervals from the full dose response model. The dashed line with a y-intercept of 0 represents where there is zero selection - positive or negative - for the plasmids. Lines of best fit and shaded bands show the median and 95% credible intervals of predictions from a model looking at how the selection rate constant changes across different combinations of temperatures and antibiotics.

**Figure S16.**
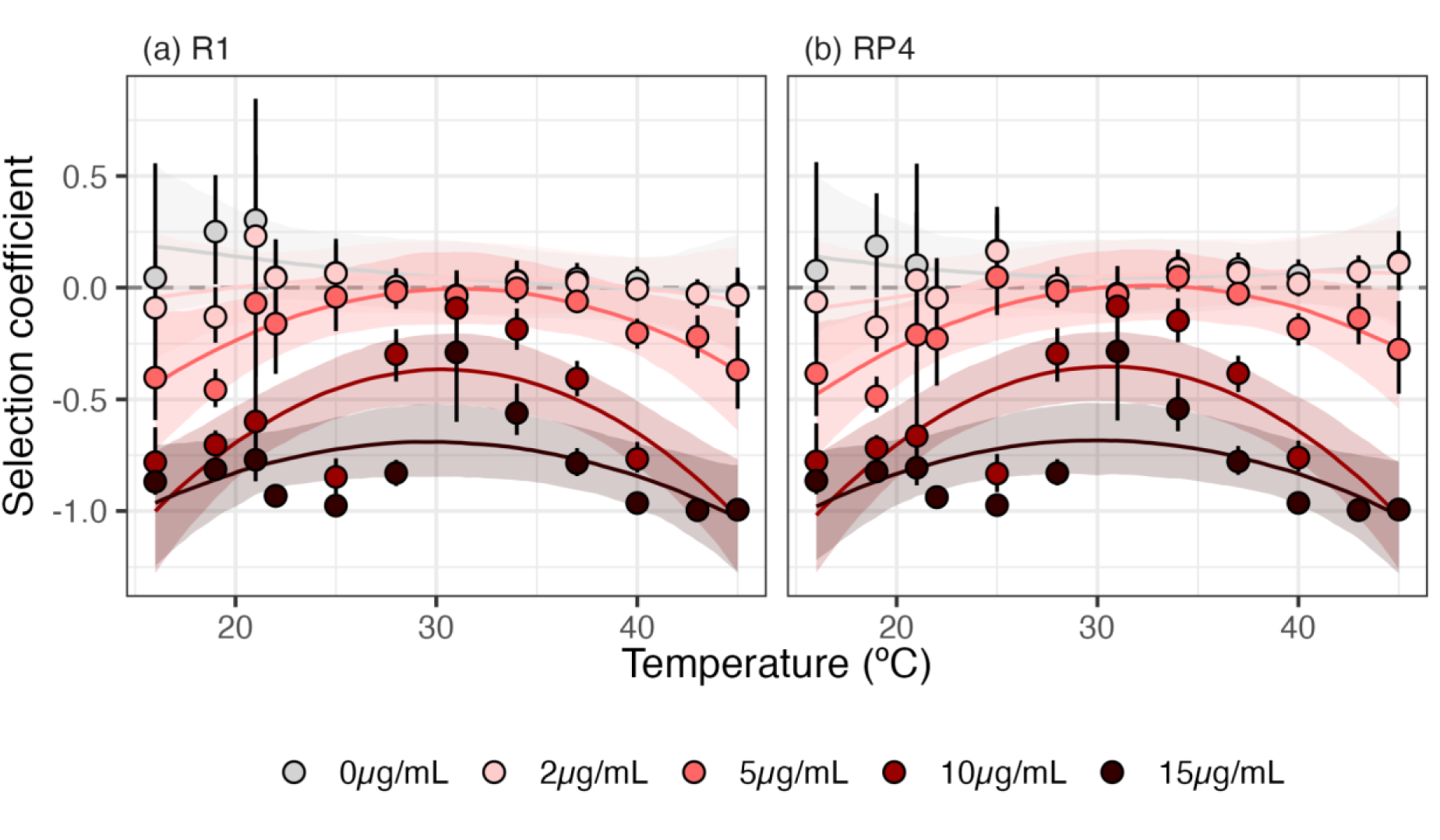
Selection coefficient. After normalising for the impact of temperature, selection for resistance is strongest at low and high temperatures. Negative values reflect selection for the plasmid, and positive values reflect selection against. Selection changes non-linearly with changes in antibiotic concentration and temperature. At concentrations close or lower than the smallest NSEC, we see no selection for resistance (all predictions have credible intervals crossing 0). At 5µg/mL, we see selection for AMR from 16-23°C and beyond ∼40°C for both plasmids, but no significant selection at intermediate temperatures. At higher concentrations, we see selection at all temperatures. Points and error bars represent median estimates and 95% credible intervals from the full dose response model. The dashed line with a y-intercept of 0 represents where there is zero selection - positive or negative - for the plasmids. Lines of best fit and shaded bands show the median and 95% credible intervals of predictions from a model looking at how the selection coefficient changes across different combinations of temperatures and antibiotics.

**Figure S17.**
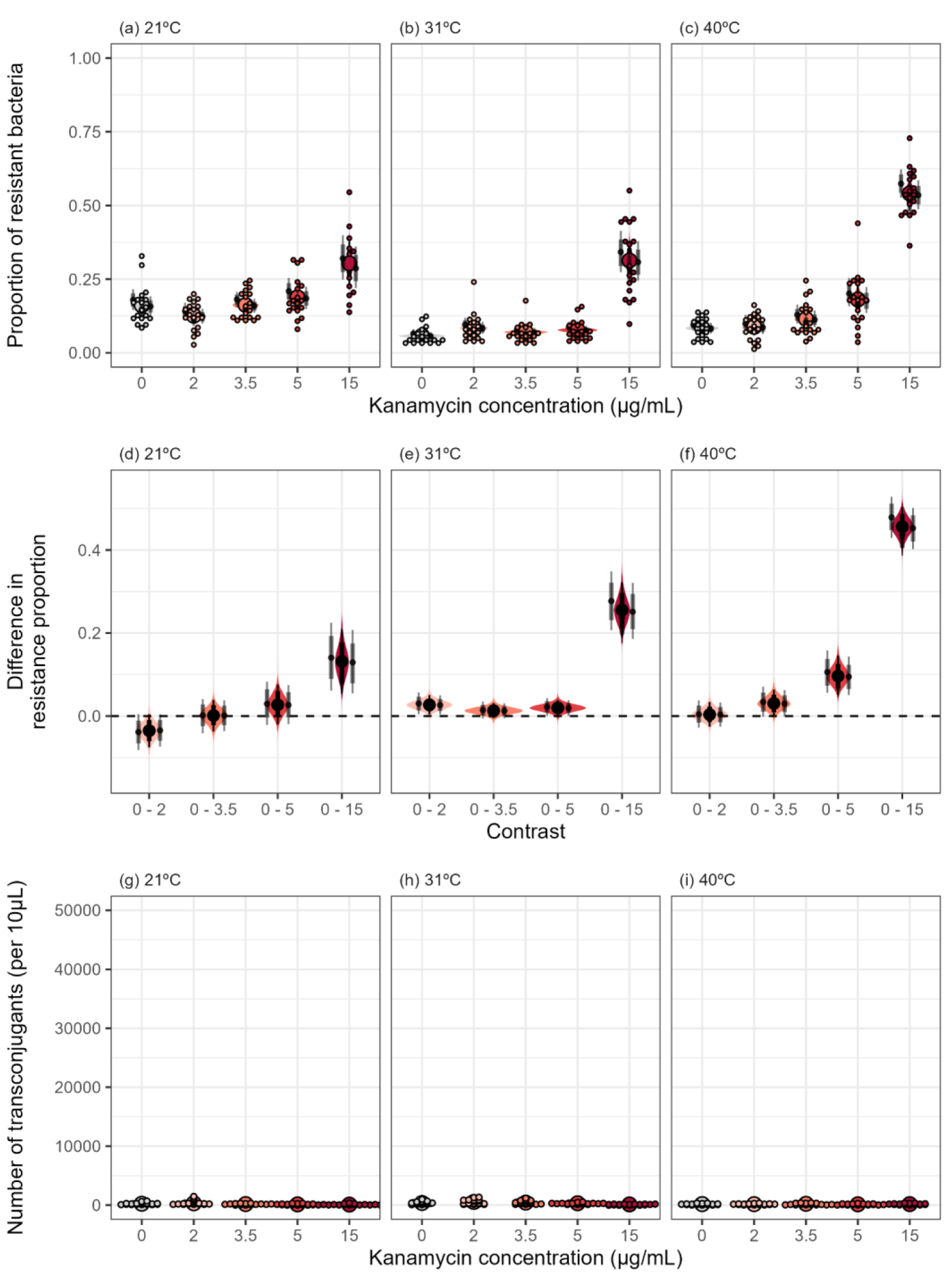
Temperature- and kanamycin-dependent resistance invasion and RP4 plasmid transfer in mixed populations. Estimates of the proportion of resistant bacteria across different kanamycin concentrations at (a) 21°C (b) 31°C and (c) 40°C. (d-f) The difference in resistance proportion measured at different kanamycin concentrations compared to zero antibiotics across temperatures. (g-i) Plasmid transfer across temperatures and kanamycin concentrations. Large coloured points and error bars represent marginal median estimates and 95% credible intervals from the models, with shaded violin bands representing the posterior distribution. Small black points and error bars represent median estimates and 95% credible intervals of the replicate level estimates. Small individual points represent individual data points.

## Notes

https://github.com/padpadpadpad/ecoli_temperature_amr

### Competing Interest Statement

The authors have declared no competing interest.

